# Unwelcome guests: characterizing the ecological niche of insertion sequences within prokaryotic genomes

**DOI:** 10.1101/2025.07.22.666184

**Authors:** Flora Gaudillière-Jami, Sophie Abby, Thomas Hindré, Ivan Junier

## Abstract

Insertion sequences (ISs) are widespread prokaryotic transposable elements, often regarded as genomic parasites that primarily cause deleterious mutations. However, they can also promote adaptive changes. These antagonistic properties make their overall impact on prokaryotic evolution difficult to grasp. Here, we address this challenge by leveraging the framework of transposon ecology to analyze IS occurrences across and within 30,499 prokaryotic genomes. Combining phylogenomics with multi-scale genomic analysis, quantitative ecology, and mathematical modeling, we provide evidence that although genomes generally provide sufficient resources for IS coexistence, universal mechanisms shape their occurrence and chromosomal distribution across genomes. These include: (i) the preferential localization of ISs within highly variable and GC-heterogeneous chromosomal regions of genomic plasticity (RGPs), which act as the primary reservoir of IS niches; (ii) a linear scaling between IS abundance and niche size, with an average of 5.4 additional accessible insertion sites per IS; (iii) a dependence of IS occurrence on the presence of other ISs, suggesting a form of group behavior; (iv) the accumulation of AT-rich sequences in both coding and non-coding regions up to 100 kb surrounding ISs, indicative of ecological isolation; and (v) the spatial partitioning of mobile genetic elements around ISs, reminiscent of ecological niche differentiation. Besides these general principles, we also uncover niche specificities associated with particular IS families, hinting at regulatory mechanisms that modulate IS activity. Altogether, this comprehensive transposon ecology approach offers new insights and avenues for understanding IS-host interactions and genome evolution, moving beyond traditional host-centric perspectives.

## Introduction

Insertion sequences (ISs) are the smallest transposable elements (TEs) found in the majority of prokaryotic genomes (1; 2). They can move within a genome by either copy-paste or cut-and-paste transposition, and can be transferred between hosts via intercellular mobile genetic elements (MGEs), such as plasmids. Upon transposition, they generate structural variations within genomes, making them key contributors to genomic plasticity and evolvability (3). They can insert into genes and disrupt their expression (2) as well as cause large-scale genomic deletions and inversions (4). These genomic changes are often deleterious to host fitness, which is why ISs have often been viewed as genomic parasites. However, they can occasionally generate beneficial mutations, for instance by knocking out costly genes (5), modulating functional elements such as gene promoters (6; 7), or facilitating the horizontal gene transfer (HGT) of antibiotic resistance genes (8).

These antagonistic properties of ISs – being often detrimental yet sometimes offering evolutionary advantages – combined with their highly dynamic nature, make their overall impact on prokaryotic evolution and genome biology difficult to grasp. Consequently, the frequent occurrence of ISs within chromosomes (9) raises questions about the factors influencing their genomic location and maintenance (1). Several of these factors have already been reported. At the local scale, several ISs are known to recognize specific DNA motifs (10; 11), whereas others show sensitivity to the AT richness of the targeted regions (1). At the chromosome scale, a mutation accumulation study in *Escherichia coli* has shown that ISs tend to insert preferentially into non-coding regions (11). This study also reported that insertions often occur near preexisting copies of the same family. Additionally, a comparative study of 80 bacterial species revealed that about 20% of IS copies are located in chromosomal hotspots representing a small fraction (1%) of chromosomes and corresponding to recent HGT (12).

Altogether, these observations raise fundamental, unresolved questions, including: i) How can we explain the widespread presence of ISs across prokaryotic genomes? ii) Which genomic features of chromosomes constrain the evolutionary maintenance of ISs? iii) To what extent does this maintenance depend on the characteristics of the IS itself? To tackle these questions, a promising avenue comes from the field of transposon ecology (13), where TEs are viewed as independent entities inhabiting host genomes and potentially interacting through mechanisms such as competition or cooperation (14; 15; 16; 17). TE-centric ecological approaches have been applied to eukaryotes, offering novel insights into the distribution and abundance of eukaryotic TEs (16; 18; 19). In prokaryotes, the ability of ISs to self-replicate, their interactions with other MGEs, and the diversity of host-IS relationships make them ideal candidates for these approaches (1; 15; 16). However, despite growing recognition, ecological approaches applied to ISs have remained largely anecdotal.

Here, we fill this gap and demonstrate how a comprehensive transposon ecology approach can provide novel, quantitative insights into the determinants of IS occurrence both across and within prokaryotic genomes. To this end, we treat each IS as an individual belonging to a specific species (the IS family) and interacting with its ecosystem (the host cell), whose characteristics – such as genomic GC content, gene and MGE density, and specific sequence features – can influence its ability to transpose and survive at specific loci. A central concept in ecology concerns the *ecological niche* of a species, which broadly represents the space it occupies within an ecosystem (20). By analogy with the environmental niche, defined as the set of conditions that permit a species’ long-term existence, here we define the niche of an IS as the genome regions and the properties of the underlying DNA sequences associated with its persistence.

Using this framework, we address the following questions, difficult to formulate within a classical host-centric genomics approach: How diverse is the IS species community within a given genome, and what determines it? What are the properties of the ecological niches of ISs within genomes? Do all IS families share the same ecological niche, or is there evidence of niche specificity? To that end, we leverage the large collection of prokaryotic genome sequences available in public databases, along with automated tools for IS annotation (21; 22; 23), analyzing the distribution of IS elements from 30 IS families across and within 30,499 complete prokaryotic genomes.

## Results

### Transposon ecology terminology

Ecological concepts and quantities for transposons have mainly been discussed and analyzed in eukaryotes. Drawing on previous work (15; 24), we therefore begin by clarifying the ecological concepts associated with prokaryotic ISs, redefining them in a genomic context:

- **Individual**: IS copy.
- **Species**: IS family as defined in the curated ISFinder database. In the ecological context, a family is understood as a lineage of ISs that share specific biochemical properties (e.g., similar transposition mechanisms, target preferences) and sequence similarity.
- **Species richness**: number of distinct IS families within a genome.
- **Population**: set of IS copies within a given genome that belong to the same family.
- **Community**: set of IS copies within a given genome, regardless of family.
- **Ecosystem**: host cell, in which an IS copy can interact with various elements including DNA sequences (e.g., host genes or other MGEs), proteins (such as host factors involved in DNA metabolism), and RNA molecules.
- **Ecological niche**: in ecology, the niche corresponds to the environmental conditions that allow the persistence of a given species. For ISs, we define the ecological niche as the range of genomic environments in which an IS can insert and persist without being lost or causing the death of the host cell. In particular, IS persistence at a locus might be influenced by a variety of genomic factors, including the function, proximity, and orientation of neighboring genes, the level of IS clustering within a genome, as well as genomic characteristics such as GC content or secondary structures.

### IS communities within genomes are well-balanced and diverse

Observing that 85% of genomes with at least one IS contain more than one IS family (Fig. 1A), and that multiple IS families often coexist within a given genome (Fig. 1B), we first asked whether IS families tend to coexist in similar population sizes, or whether competition leads, for instance, to the dominance of a single, fitter family. To address this, we computed the evenness of family composition (i.e., of IS species richness) within genomes using a descriptor commonly applied in ecology: the rank abundance curve (RAC). This curve provides key yet easily interpretable information about community composition by showing the proportion of species (IS families) as a function of their decreasing rank in abundance within ecosystems (host cells) (25; 26).

**Figure 1.**
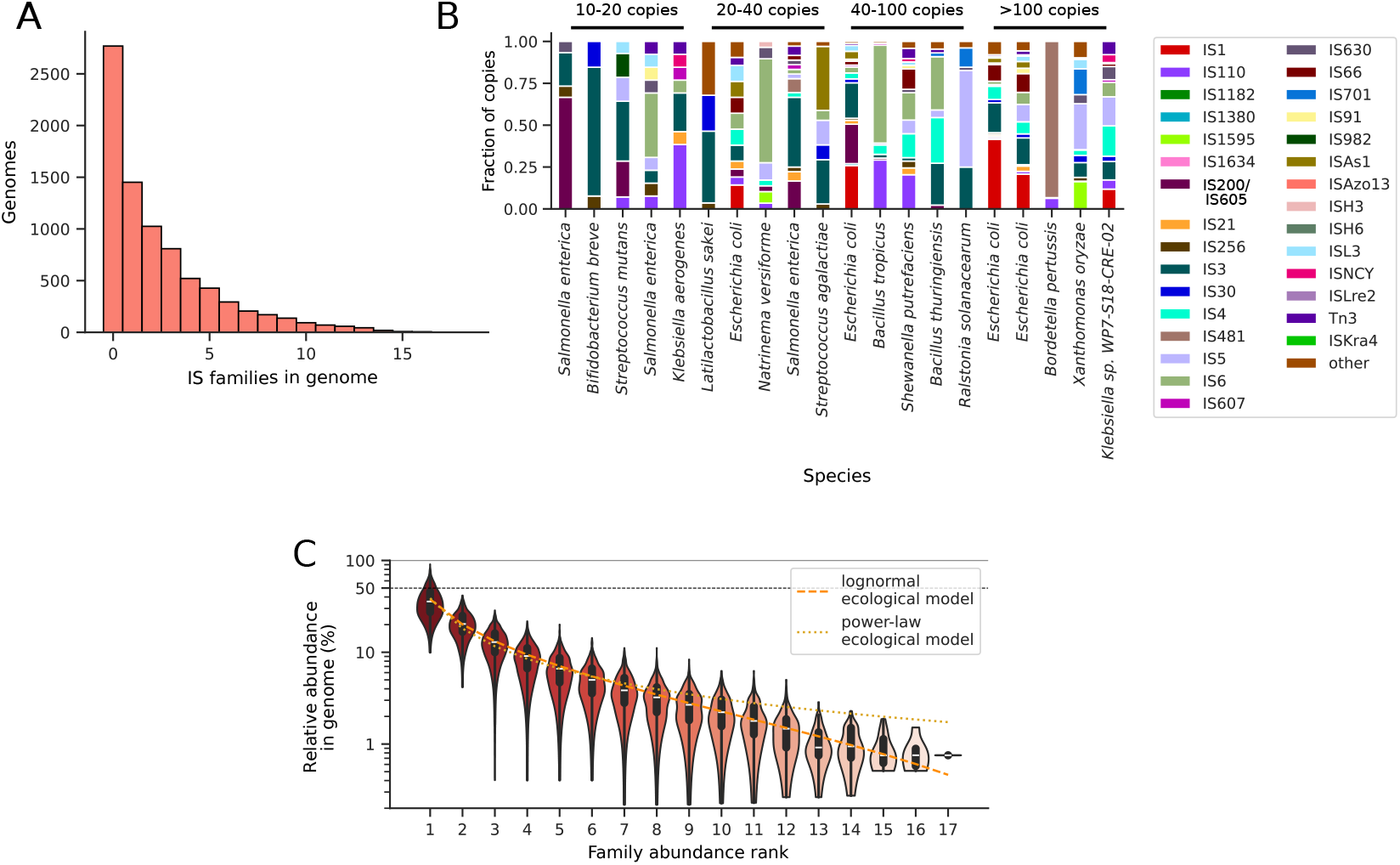
Diversity of IS communities within genomes. A) Histogram of the number of distinct IS families in prokaryotic genomes. B) IS content of 20 sample genomes from the dataset with total number of IS ranging from 10-20 to more than 100 copies. The label “other” refers to ISs that are not clearly attributed to a specific family (Methods). C) Best fits of the rank abundance curve of ISs using a lognormal and a power law ecological model.

In particular, RACs derived from models or empirical data often fall into a few distinct shapes – typically lognormal, logseries, or power law – each reflecting different levels of evenness in species composition (27). Our analysis for the ISs shows a RAC that is particularly well-fitted by the lognormal model (Fig. 1C), and, for instance, much less so by the power law model (Fig. 1C). This suggests a well-balanced (25; 27), mosaic nature of IS communities within prokaryotic genomes, where no single IS family dominates excessively and families rarely contribute only a small fraction of copies. In other words, provided ISs can enter a genome, the latter offers sufficient niche resources to support their coexistence.

### IS niche availability is strongly constrained by the functional organization of genomes

We investigated the local properties of the ecological niches occupied by ISs within host genomes by examining their immediate genomic contexts. Corroborating results from mutation accumulation lines in *E. coli* (11) and extending them across species, we find that ISs preferentially locate in intergenic regions rather than in regions where they interrupt coding sequences (CDSs) (Fig. 2A). This trend holds at the family level for 29 of the 30 IS families, all of which show a significant signal (Fig. S1). We then asked whether these intergenic regions are linked to specific aspects of the functional organization of genomes, which is known to be highly constrained (28; 29). In particular, genes often belong to operons, and cofunctional operons are themselves organized into larger genomic units that tend to remain in close proximity throughout evolution, a property known as gene synteny. Analyzing *E. coli* genomes for which accurate functional genome annotations are available, and which harbor 19 of the 30 known IS families, we find that identifiable intergenic ISs are disproportionately located in inter-operonic regions (Fig. 2B), with this pattern holding for all but one IS family (Fig. S2). Furthermore, inter-operonic ISs rarely reside in-between syntenic gene pairs (Fig. 2C), with the notable exception of the IS*30*, IS*66*, and IS*91* families, which exhibit a significant opposite trend (Fig. S3).

**Figure 2.**
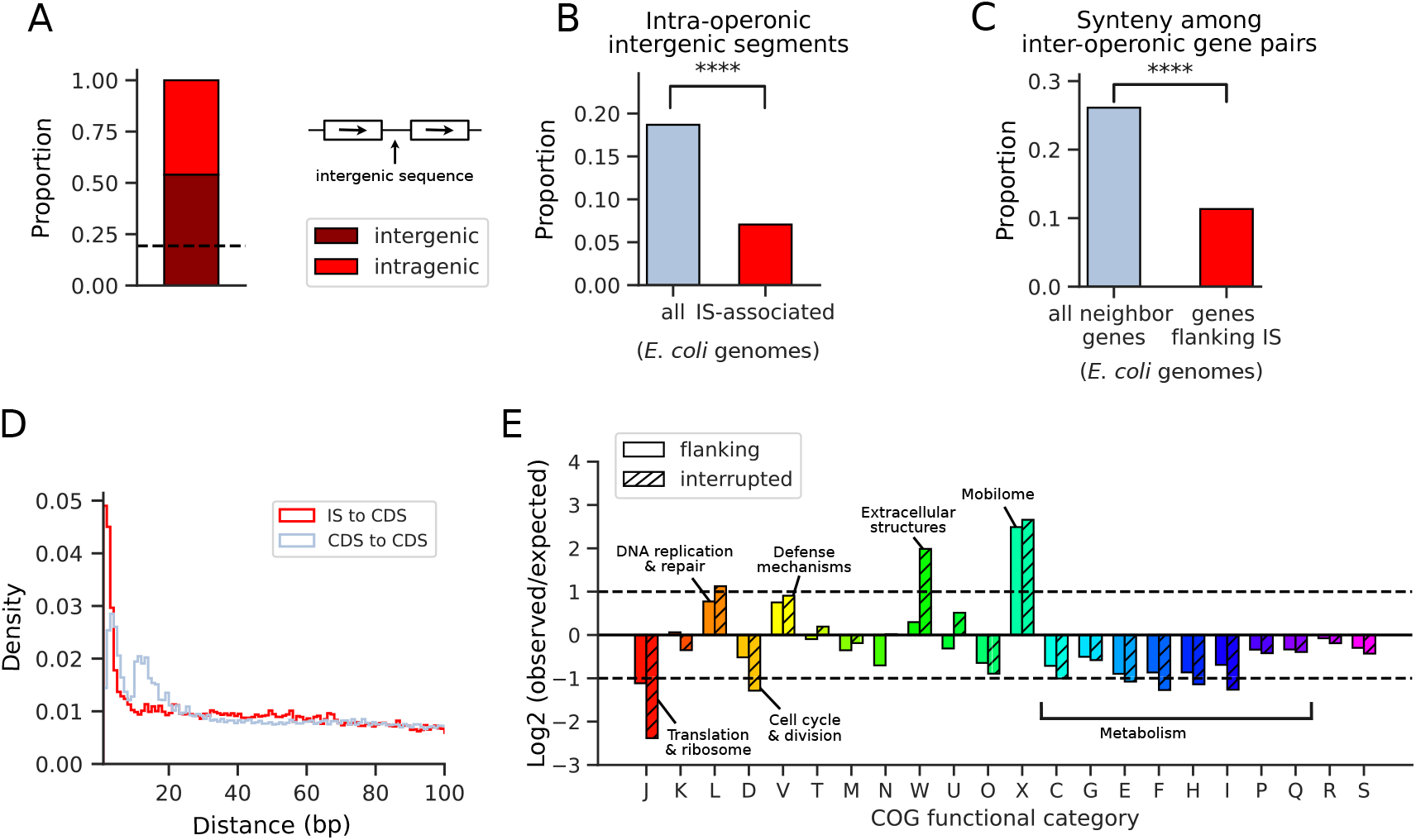
Functional environment of IS elements. A) Proportion of identifiable IS transposition sites found in intergenic segments (dark red) and intragenic segments (red). Dotted line: average proportion of intergenic sequences in prokaryotic genomes. B) In *E. coli* genomes, proportion of intergenic segments found inside of an operon (light blue) and proportion of intergenic ISs found inside of an operon (red). The statistical significance (*P <* 0.001) was computed using a chi2 test. C) In *E. coli* genomes, proportion of syntenic gene pairs among consecutive genes that belong to different operons (light blue) and proportion of syntenic gene pairs among genes flanking ISs that are found neighbor in at least one genome (red). The statistical significance (*P <* 0.001) was computed using a chi2 test. D) Normalized distribution of distances from ISs to their flanking CDSs (red) or between random CDSs (light blue), for distances ranging from 1 bp to 100 bp. E) Functional enrichment of genes flanking ISs (plain) or interrupted by ISs (striped). Enrichment or depletion is measured by calculating the log 2[*f*_*obs*_*/f*_*exp*_], with *f*_*obs*_ the frequency at which a given functional category is found in either genes flanking ISs or interrupted by them, and *f*_*exp*_ the frequency at which this functional category is found in randomly sampled genes. Horizontal lines indicate 2-fold increase and decrease, respectively. Not indicated COG annotation: K: transcription; T: signal transduction; M: cell wall/membrane; N: cell motility; U: trafficking & secretion; O: protein modification; C: energy production; G: carbohydrate metabolism; E: amino acid metabolism; F: nucleotide metabolism; H: coenzyme metabolism; I: lipid metabolism; P: ion transport; Q: secondary metabolism; R: general function prediction only; S: unknown function.

The analysis of the location of intergenic ISs with respect to their flanking genes reveals two opposite trends. First, CDSs tend to be located farther from ISs than from random CDSs (Fig. S4A, Fig. S5). In contrast, when focusing on the occurrence of short distances below 100 bp, we find an over-representation of IS insertions occurring just one, two, or three base pairs upstream or downstream of CDSs (Fig. 2D), a pattern observed in 28 of the 30 IS families (Fig. S4C, Fig. S6). Finally, and consistent with recent observations (9), we observe that 17% and 50% of IS families are preferentially found between divergent and convergent gene pairs, respectively (Fig. S4B).

We examined in more detail the functions of genes interrupted by, or flanking, ISs by assigning all genes in our dataset to one of 22 bacterial Cluster of Orthologous Genes (COG) functional classes (Fig. 2E). Similar patterns are observed for interrupted and flanking genes, though functional biases are systematically stronger for interrupted genes – the only exception is a strong over-representation of extracellular structure functions (class W) observed only among interrupted genes; closer inspection reveals that this is largely driven by the recurrent disruption of a single gene, *fimC* involved in pilus assembly of type I fimbriae. Specifically, we observe that metabolism-associated functions globally tend to be under-represented, whereas some specific functions are strongly over-represented. These include mobilome genes (X), consistent with previously reported MGE hotspots (12), as well as defense mechanisms (V) and DNA replication and repair (L). The latter is particularly intriguing given the essentiality of these functions – for example, translation (J), and cell cycle (D) are strongly under-represented.

Altogether, these results reveal a strong influence of the functional organization of genomes on IS niche availability. Much of this likely reflects purifying selection, as ISs tend to be under-represented in regions under selective pressure. However, the over-representation of specific functions among IS-interrupted genes such as defense mechanisms or DNA replication and repair may instead suggest positive selection, either mediating host adaptation to new environments (30) or stress conditions (31), or possibly favoring the ISs themselves. Moreover, recurrent patterns in the local organization of ISs with respect to their flanking genes suggest that ISs often play specific roles in modulating the expression of nearby genes, as has been demonstrated experimentally in specific cases (32).

### Terminator-like sequences are over-represented upstream of ISs

We observe a strong over-representation of DNA sequences characteristic of intrinsic transcription terminators located immediately upstream of ISs, in sharp contrast to the typical tendency of intrinsic terminators to be found downstream of genes (Fig. 3, Fig. S7) – the strongest signal is observed for the IS*200*/IS*605* family, for which experimental evidence for the activity of such terminators already exists (33). Furthermore, in some IS families, terminator-like sequences can also be found inside of IS elements (between the limits of the IS and the transposase CDS, Fig. S8). Overall, we thus find that several IS families are disproportionately found in genomic contexts likely to limit transposase expression, which we interpret as a characteristic that promotes their persistence within genomes.

**Figure 3.**
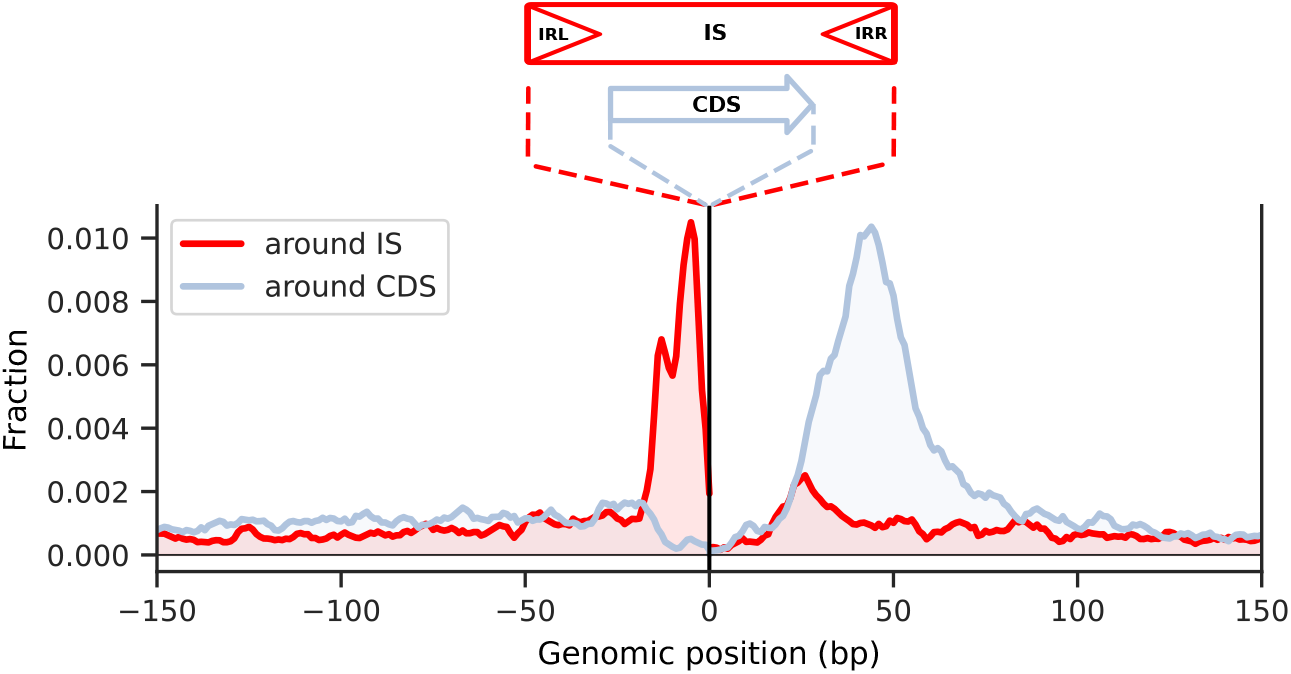
Genomic distribution of terminator-like sequences around ISs. Frequency of terminator-like sequences in the 150 bp upstream and downstream of ISs compared to the same frequency in the vicinity of random CDSs. Genomic positions are shown relative to the orientation of CDSs. The detailed signal associated with each family is provided in Fig. S7. Note that while most IS families are flanked by inverted repeats (IR) as depicted here, some families such as the IS*200*/IS*605* are not.

### Clustering properties: IS niche size is predicted to scale with IS number

To better understand the behavior of ISs along chromosomes and to apprehend the nature of their niche, we investigated whether ISs tend to colocalize. To this end, we analyzed the tendency of ISs to cluster, considering ISs to be part of the same cluster if they are adjacent along the DNA and separated by less than 2.5 kb (Methods). In agreement with (12), we observe that in 19% of cases, an IS is part of a cluster, with cluster sizes reaching up to 18 individuals and showing diverse compositions regardless of size (Fig. S9A). The composition of a given cluster strongly varies across strains of the same species, as can be seen in *E. coli* (Fig. 4A). Therefore, chromosomal IS clusters are likely to result from a dynamical, stochastic birth-and-death-like process (34) occurring at those loci, rather than from en bloc integration of plasmid-carried IS clusters. To better apprehend this underlying process, we computed the statistical distribution of cluster sizes. We observe that for chromosomes with fewer than 120 ISs, which corresponds to 96% of our dataset, the distribution follows an exponential law with a decay constant that is independent of both the total number of IS copies (Fig. 4B) and overall IS density (Fig. S9B) – we also verified that the same exponential distribution of cluster size holds across various phylogenetically distant species (Fig. S10). In chromosomes with both more than 120 ISs and high IS density, the distribution deviates from an exponential, becoming heavier-tailed (Fig. S9C). Such a “universal” (exponential) law (35) in chromosomes containing fewer than 120 ISs further supports the existence of a generic stochastic process driving the chromosomal location of ISs, independent of their features. To characterize this process, we considered three models of random locations of ISs along the chromosome, each representing a different hypothesis regarding the accessibility of genomic regions by the ISs : (I) the entire chromosome is accessible to ISs, (II) accessible regions are restricted to a fraction of the chromosome, and (III) the number of accessible sites increases proportionally with the total number of ISs – details of the models and their predictions are provided in Supp. Note 1. We find that the model (III) with proportional accessible sites, and only this model (Supp. Note 1), leads to an exponential distribution that is independent of the number of ISs in the genome. Fitting the data yields an estimated number of accessible sites equal to approximately 5.4 times the number of ISs per chromosome (Supp. Note 1). Consistently, only this model can account for the observed dependence of the mean cluster size per chromosome as a function of IS counts, up to 120 ISs (Fig. 4C). Beyond this threshold, data are well explained by assuming, instead, that the number of accessible sites saturates (i.e., scenario II) at the value predicted for 120 ISs under scenario III, that is, approximately 5.4 ×120 ≃648 IS-equivalent sites (Fig. 4C, Supp. Note 1). More precisely, the best-fitting model is one in which the size of the accessible region is proportional to chromosome size (see Supp.Note1).

**Figure 4.**
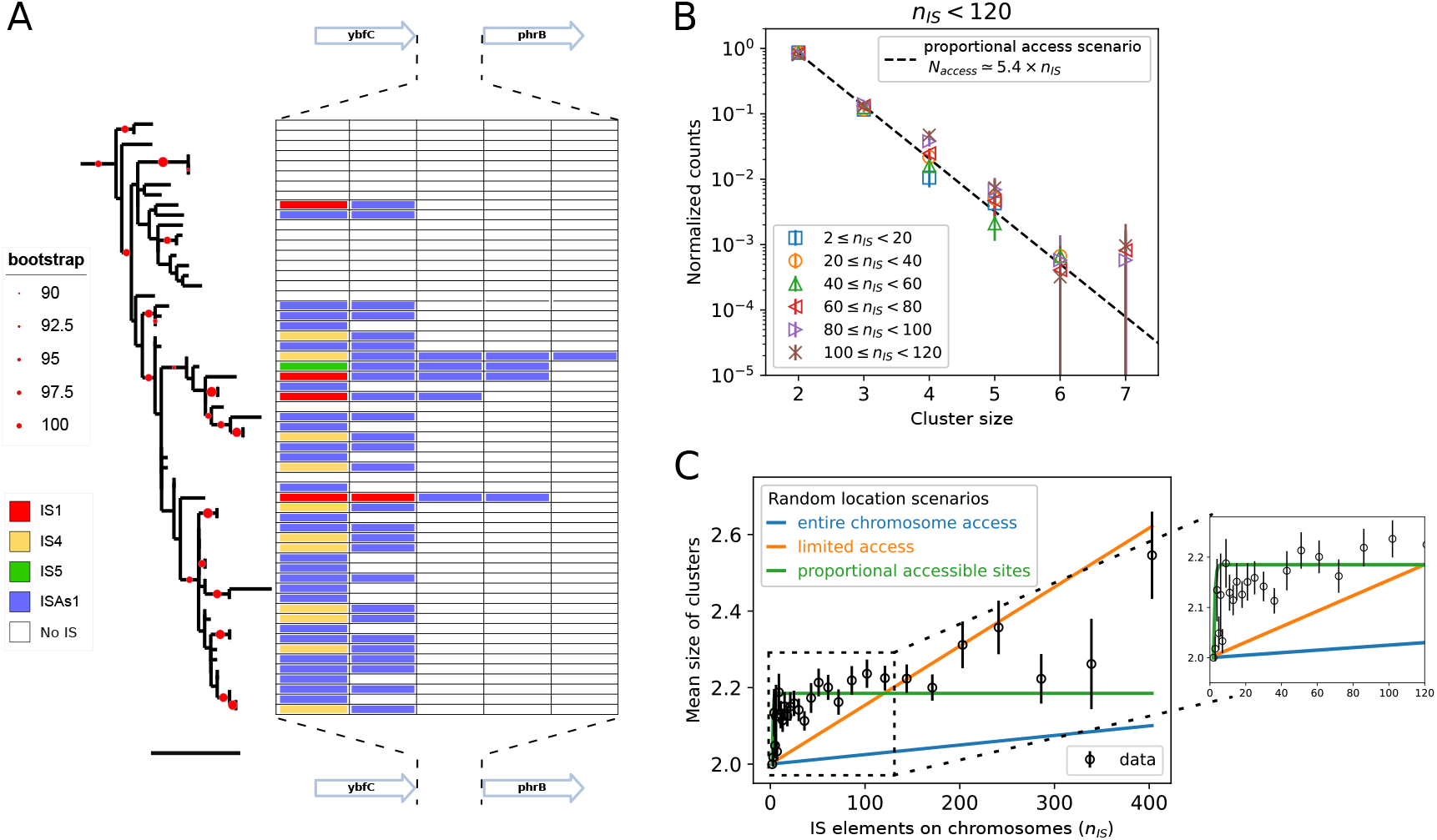
IS clusters: diversity, statistical properties and modeling. A) Example of cluster composition diversity between two conserved genes in different *E. coli* genomes. For each genome, the IS content between the two neighboring genes *ybfC* and *phrB* is represented, ranging from 0 (empty rows) to 5 elements (fully colored row). Tree scale: 0.002 substitutions per site. B) Distribution of cluster sizes in genomes containing less than 120 ISs. Each plot corresponds to a distribution that has been computed by constraining the number of ISs within the genome to belong to a certain interval. The black dashed line indicates the prediction (an exponential law) of the model of proportional accessible sites, considering that each IS is associated with approximately 5.4 accessible sites. C) Mean size of clusters as a function of the number of ISs in the corresponding chromosomes. We also report the results of three models based on random IS transpositions along the chromosome under specific constraints: i) full accessibility, where ISs can insert and survive anywhere along the chromosome (blue); ii) limited access, where ISs can only survive in a fixed portion of the chromosome (see Supp. Note 1 for more complex scenarios) (orange); iii) proportional accessible sites, where the presence of an IS is associated with additional opportunities for other ISs to transpose and survive (green). For the latter scenario, the parameters are those of the model used in panel (A). We also note that part of the residual variation around the prediction of the fixed-limit scenario (ii) can be explained by an alternative scenario, in which the limited access region is not fixed and independent of chromosome size, but instead scales proportionally with it. (see Supp. Note 1). *n*_*IS*_ : number of ISs on the chromosome.

Altogether, these results suggest that cluster formation primarily arises from limited chromosome accessibility, and that the size of the niche (the number of accessible sites) is proportional to the number of ISs.

### Regions of genomic plasticity constitute the primary reservoir for IS niches

To further characterize IS niches, we compared the distribution of ISs along chromosomes with the distribution of conserved versus variable genomic regions. To this end, for several species, we first aligned, normalized, and segmented the chromosomes of all their genomes into 1,000 bins. We then computed the occurrence profiles of ISs, persistent (or core) genes, and regions of genomic plasticity (RGPs) (36), the latter mostly containing accessory genes. In *E. coli*, we observe a strong correlation of IS locations with RGPs, along with a negative correlation with persistent genes (Fig. 5A, see Fig. S11AC for species with significantly lower and higher GC content). More precisely, 66% of ISs are located within RGPs, a proportion that reaches 87% when considering only clustered ISs. The average number of ISs in RGPs is found to scale with RGP size, often in a linear manner (Fig. 5B and Fig. S11BE) – for comparison, there exists no clear relationship between the number of ISs and the size of chromosomes (21; 9) (Fig. S14).

**Figure 5.**
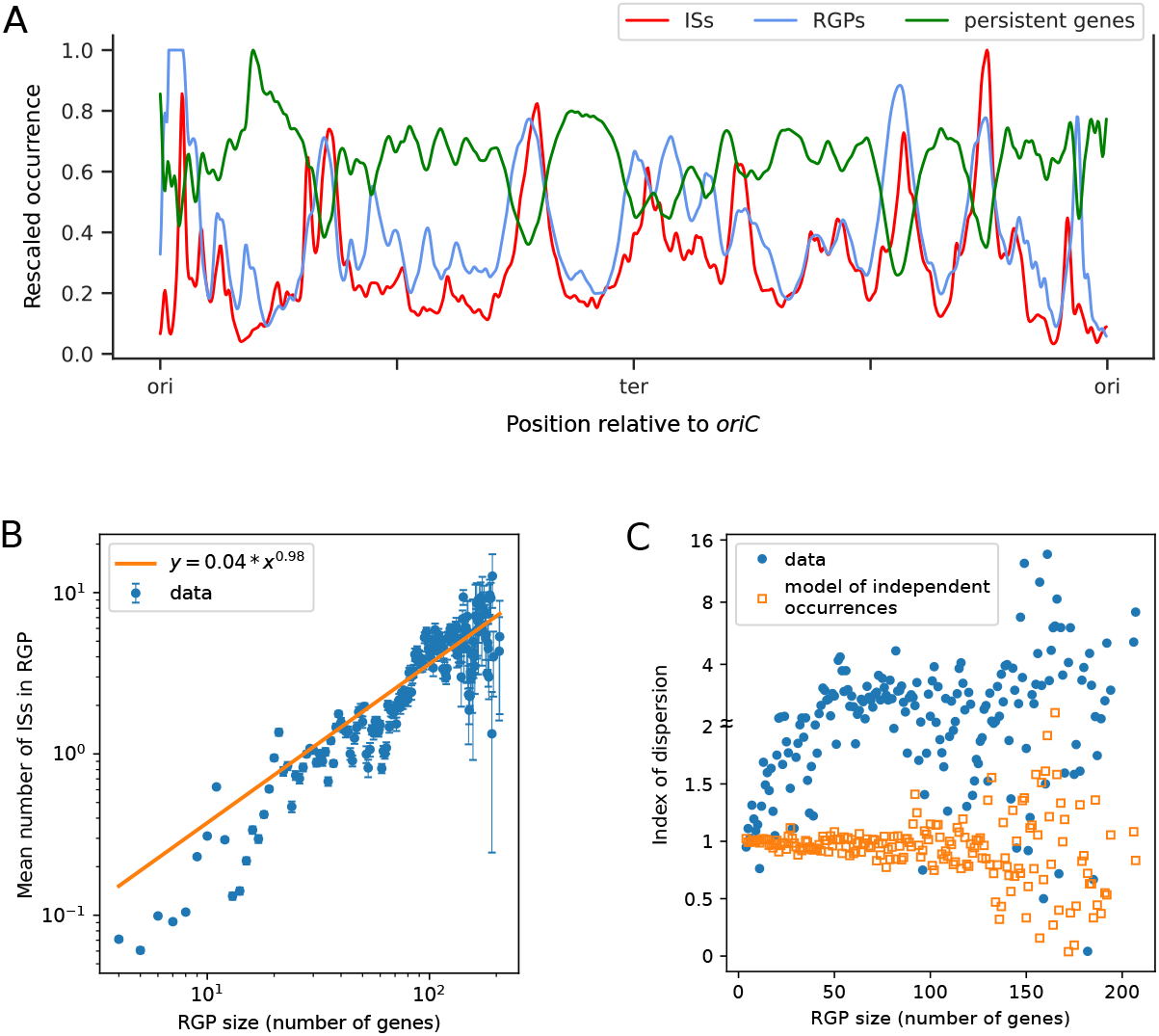
ISs versus RGPs in *E. coli* (see Fig S11 for other species). A) Distribution of unique IS positions (red), persistent gene positions (green) and frequency at which a region is a region of genomic plasticity (RGP, blue) across *E. coli* chromosomes. For each profile, values were rescaled to the maximum value within that profile. B) Mean number of IS copies as a function of RGP size (number of genes within each RGP), computed from all RGPs found in *E. coli* chromosomes, using a log-log scale – error bars represent the standard error of the mean. The orange line indicates the best power-law fit, revealing an almost linear relationship. C) Index of dispersion (i.e., ratio of the variance to the mean) for the IS content of RGPs as a function of RGP size. The orange symbols represent values obtained in a model in which, within a given RGP, genes have a fixed probability of being an IS, as determined by the fit in (B) (approximately 1*/*28 here), independent of other IS occurrences. Note that the y-axis uses a semi-logarithmic scale to accommodate points with very large values, with a linear scale up to 2 (indicated by the *≈*) and a logarithmic scale above.

Altogether, these results suggest that RGPs serve as the primary reservoir for IS niches, and that ISs often contribute, in average, to a constant proportion of each RGP in a given species.

### IS occurrences in RGPs are not independent from each other

We investigated whether ISs behave independently within RGPs or if there is evidence of interactions suggestive – from an ecological perspective – of group behavior. To address this, we measured the dispersion of IS copy numbers within RGPs by calculating the variance-to-mean ratio, a metric known as the index of dispersion in ecological studies. Under the assumption of independent IS insertions, this index is expected to be close to 1, regardless of RGP size – reflecting a Poisson distribution where the probability of an RGP gene being an IS is fixed. However, using the same dataset as that we used to compute the mean number of IS copies for all RGPs of a given size (see Fig. 5B), our results reveal overdispersion, with most index values exceeding 1 (Fig. 5C, Fig. S11CF). We therefore conclude that IS occurrences within RGPs are not independent of each other, but rather indicative of group behavior.

### IS niches are AT-enriched and extend over tens of kb of coding and non-coding DNA

To further investigate the genomic properties of IS ecological niches, we compared the chromosomal distribution of ISs to that of GC content. To this end, we followed the same procedure we used to compute the occurrences of RGPs and persistent genes (see above). Considering three distantly related bacterial species, we find that the IS distribution strongly correlates with the average AT content profile computed over the chromosomes, regardless of the overall GC content of the species (Fig. 6A and Fig. S13AC). This correlation is barely visible in individual genomes but becomes clear when analyzing multiple chromosomes (typically 10 or more in *E. coli*) (Fig. S12), that is, when averaging the signal over evolutionary time. In line with this, and with the preferred location of ISs within RGPs, we also find that the IS distribution strongly correlates, in a given species, with the profile of GC variance computed over its chromosomes (Fig. 6B and Fig. S13BD).

**Figure 6.**
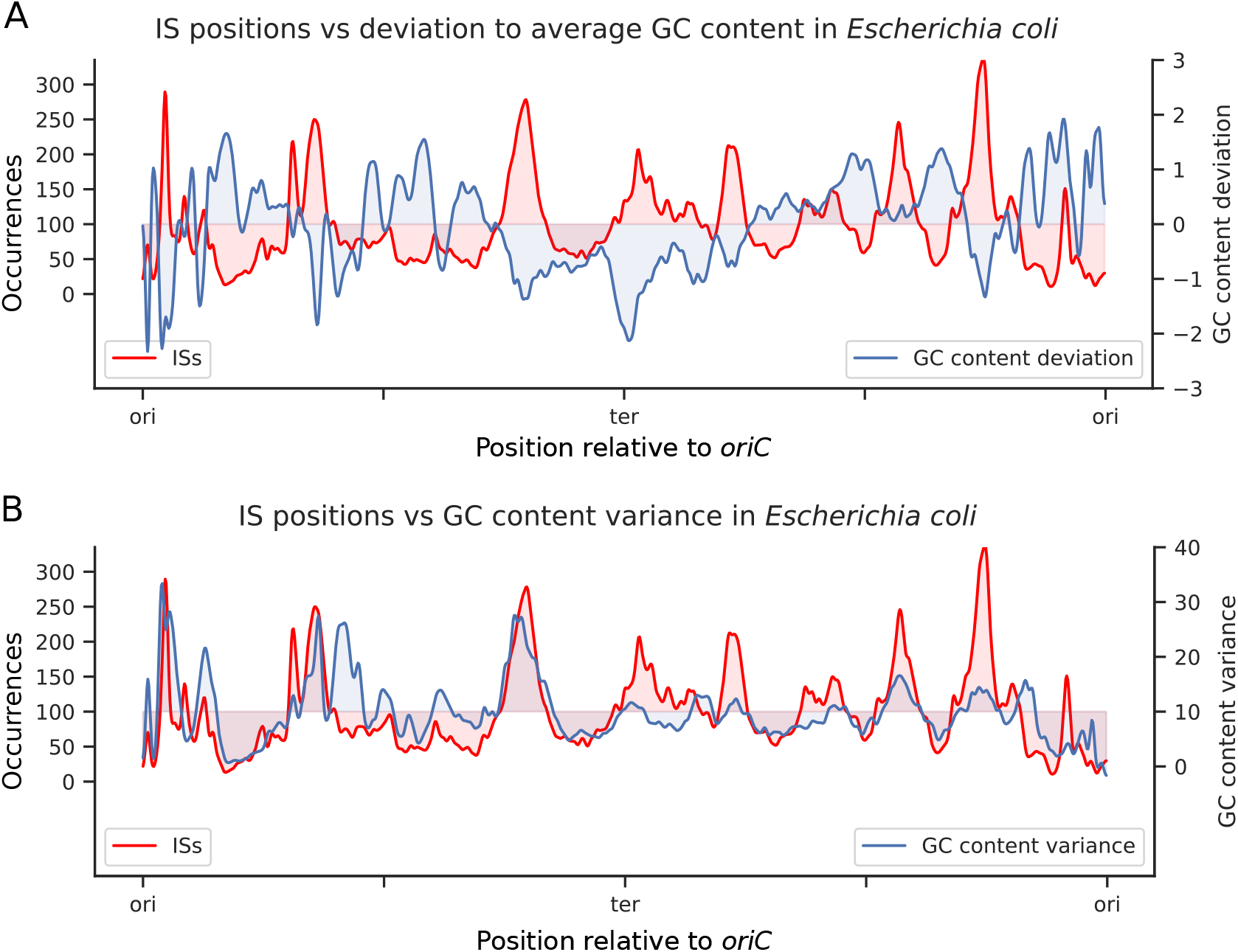
Characterization of the GC content of IS niches in *E. coli* chromosomes (see Fig S13 for other species). A) Distribution of unique IS positions (red) and deviation to average GC content (blue) across *E. coli* chromosomes. Pearson correlation: -0.47. B) Comparison of ISs distribution to the variance in GC content across *E. coli* chromosomes. Pearson correlation: 0.51.

To assess the generality of this AT enrichment in the vicinity of ISs – and given the impossibility of globally aligning all bacterial genomes – we aggregated all DNA sequences flanking ISs (accounting for IS reading direction) from our dataset of 30,499 genomes, and computed the average GC content across the corresponding pile-ups. Fig. 7A presents the resulting profile as a function of the distance from the IS borders and compares it to profiles generated by aggregating sequences around random CDSs. We find a marked GC depletion (i.e., AT enrichment) around ISs that extends over tens of kilobases. Such AT enrichment can stem from two main sources: either a depletion of CDSs (which are richer in GC compared to intergenic regions), or an intrinsic AT enrichment occurring in CDSs, intergenic regions, or both. By quantifying contributions to the signal (Supp. Note 2), we find that the observed trend arises primarily from intrinsic AT enrichment (Fig. S15), which concerns both CDSs and intergenic regions (Fig. 7B). Moreover, it affects CDSs independently of the codon position (Fig. 7C), and regardless of the GC content of genomes. The strongest effect, however, systematically concerns the codon position with the highest overall GC content (Fig. S16).

**Figure 7.**
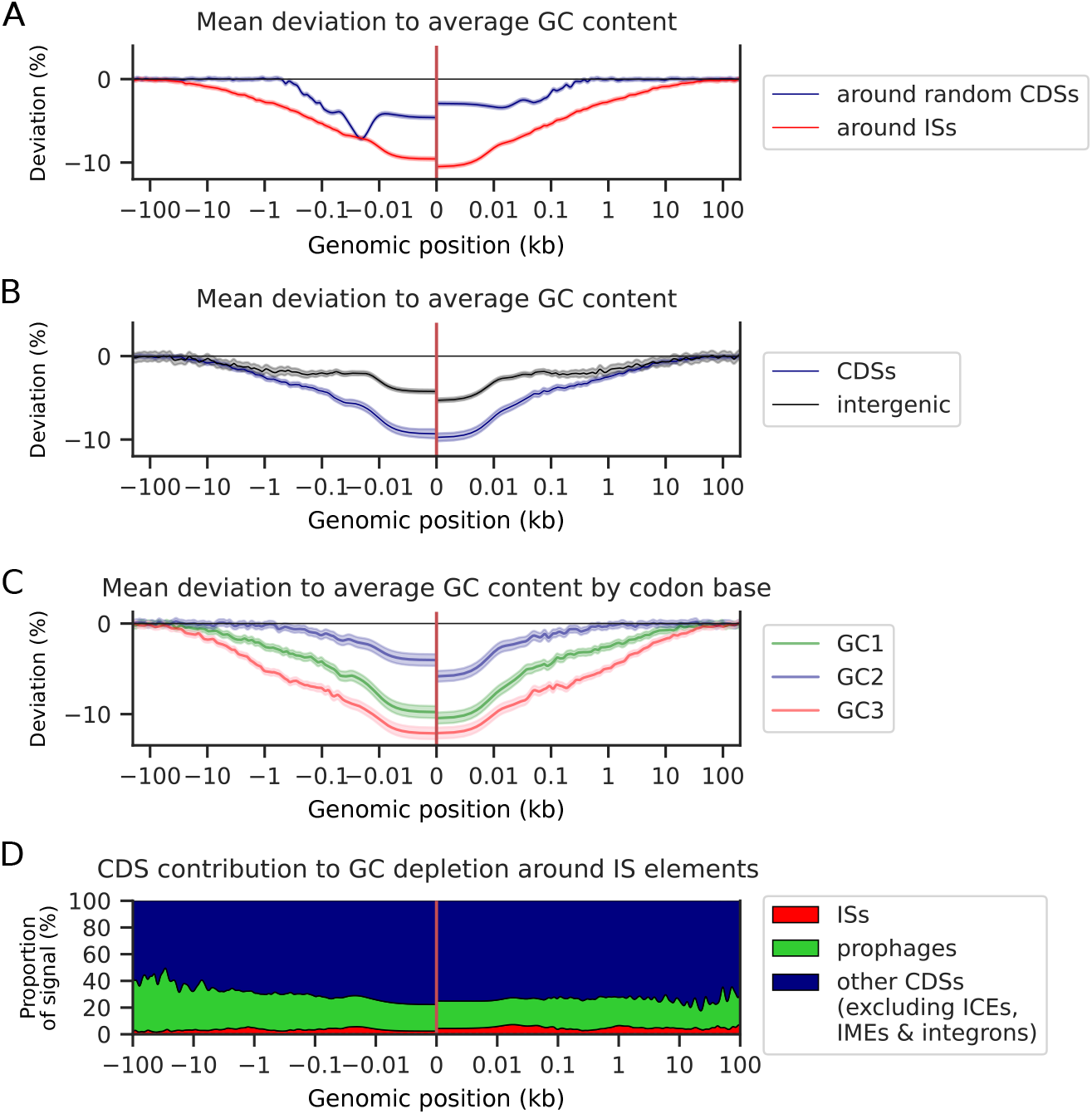
GC content deviation around ISs computed from all 30,409 genomes. A) Relative deviation (in %) from average GC content at a given distance from a CDS (blue) or an IS (red), considering its transcriptional orientation as in Fig. 3. B) Relative deviation (in %) from average GC content at a given distance from an IS, in CDSs (blue) or intergenic sequences (gray). C) Relative deviations from average GC content, shown separately for each codon position. D) Contribution of different types of CDSs (IS CDSs, prophage CDSs, ICE & IME CDSs, integron CDSs and other CDSs) to the GC depletion around ISs. The contribution of integrons, ICEs and IMEs is too small to be visible. All panels: the dashed areas (barely visible in A) show the standard errors of the mean.

Regardless of a genome’s overall GC content, ISs are thus found in chromosomal regions that accumulate more AT-rich sequences than the rest of the genome. This AT enrichment affects both coding – independently of codon position – and non-coding sequences, and it extends over regions up to ∼100 kb around ISs.

### AT enrichment primarily originates outside known MGEs and affects them

Given that plastic regions of genomes are characterized by frequent HGT (12; 36), and that the GC content of horizontally acquired genes tends to differ significantly from that of the host genome (37) (Fig. S17), we investigated the relative contribution of known MGEs to the AT enrichment of CDSs around ISs, across all prokaryotic genomes. To this end, following methods in (12), we considered four main categories: ISs, prophages, identifiable genes of ICEs/IMEs (Integrative and Conjugative Elements/Integrative Mobilizable Elements) and integrons (Methods). We find that ∼ 60% of the AT enrichment of CDSs around ISs arises from genes other than known MGEs (Fig. 7E). Indeed, the main MGE contribution comes from prophages and accounts only for ∼ 30%, whereas ISs account for less than 5%, and ICEs and integrons together for less than 0.5%. Moreover, prophage sequences tend to become increasingly AT-rich the closer they are to an IS (Fig. S18A), a trend not observed for ISs and integrons, and only weakly for ICEs/IMEs (Fig. S18B). The majority of the AT-enrichment signal around ISs is therefore not attributable to known MGEs. Moreover, some MGEs, although already more AT-rich than the rest of the genome, exhibit an even stronger AT-enrichment when located near ISs.

### Ecological niche differentiation of MGEs

An analysis of the genomic distribution of MGEs around ISs (Fig. 8) further reveals that the identifiable genes of ICEs/IMEs (Methods) tend to locate farther from ISs than prophages or integrons. Moreover, these ICE/IME genes are depleted in the immediate vicinity of ISs and they are, on average, more GC-rich than other MGEs (Fig. S19). A characteristic length of approximately 70 kb associated with prophage locations is also clearly visible (Fig. 8, in green). A similar length has been previously reported and interpreted as an upper limit for chromosomal integration of prophage segments (38; 39), which are subsequently subject to genetic degradation (38). This suggests that prophages close to ISs may primarily result from recent HGT.

**Figure 8.**
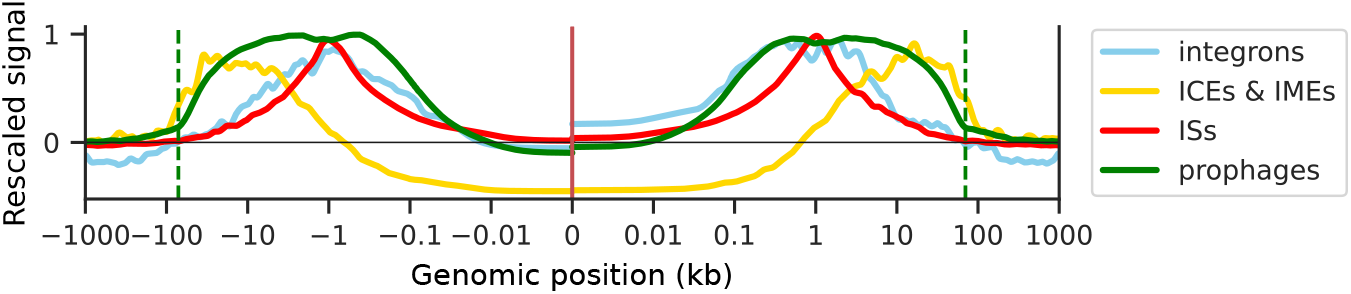
Distribution of MGEs around ISs. The signal was rescaled such that for a given MGE *M*, we compute [*f* (*M*) *− f*_*base*_(*M*)]*/*[*f*_*max*_(*M*) *− f*_*base*_(*F*)], with *f* (*M*) the fraction of sequences for which this position corresponds to the MGE, *f*_*base*_(*M*) the average fraction in the genome, and *f*_*max*_(*M*) the maximum value reached by *f* (*M*). The vertical green dashed lines indicate a characteristic length of 70 kb associated with a sharp change in the distribution of prophage locations.

Altogether, these results reveal a partitioning of the genomic space occupied by MGEs around ISs, which we interpret as evidence of ecological niche differentiation among MGEs.

## Discussion

We conducted a comprehensive ecological analysis of ISs both within and across genomes. In doing so, while keeping in mind the limitations of automated detection methods and the lack of information on the evolutionary trajectories of ISs and their genomic context (Supp. Note 3), we uncovered several phenomena that are difficult to interpret outside the framework of ecology. In particular, we found that although genomes generally provide sufficient resources for ISs to coexist, regions of genomic plasticity (RGPs), which are the most variable chromosomal regions across strains of a given species, constitute the main reservoir for IS niches. We also found that regardless of the species studied – including species with very different GC content – the size of the IS ecological niche varies linearly with the number of ISs, always maintaining the same average proportion: for each additional IS in a RGP, there are approximately 5.4 additional niche sites. These findings have two key implications. First, the observation of the same property across virtually all species suggests the existence of very generic – poorly understood mechanisms, reminiscent of the universal mechanisms underlying the scaling laws observed in the functional composition of bacterial genomes (40; 41). Second, the question of causality arises: Do ISs drive the expansion of their own niches – or of the RGPs – or do larger RGPs provide more space for ISs? The first scenario would correspond to a “niche construction” mechanism, where the insertion of an IS in a particular location increases the likelihood of subsequent IS insertions or promotes the extension of RGPs – for instance through large IS-mediated duplications of chromosomal segments, the insertion of composite transposons and mobilizable units, or the integration of IS-carrying plasmids. The second scenario would still leave open the question of what determines the frequent observation of a constant fraction of ISs within RGPs. Distinguishing these scenarios will certainly require a high amount of high-resolution temporal evolutionary data. In all cases, we find that IS occurrences within RGPs do not behave independently of each other, which could be because the presence of one IS may increase the likelihood of others inserting nearby. Rather than acting as isolated, selfish elements – a common explanation for the spread of TEs among organisms (42), – we thus propose that ISs exhibit group behavior, a hallmark of collective phenomena in microbial ecology (of organisms).

Perhaps unsurprisingly, in all the species we investigated in detail – and for which a large number of genomes were available – IS niches are found within the most dynamic chromosomal regions, the RGPs, where HGT is particularly frequent. Nevertheless, the observation of a specific often linear – scaling of IS number with RGP size suggests that the internal dynamics of these RGPs is driven by specific composition rules. In this context, knowing that RGPs can contain hundreds genes, and that IS-surrounding genomic patterns – such as the GC depletion signal or the location of MGEs – extend up to 100 kb, the HGT hotspots identified in previous studies, which typically comprise only a few percent of the genome (12), may reflect finer-scale structuring of ISs and MGEs within these RGPs. Along this line, our observation of MGEs and ISs occupying specific genomic locations relative to each other, with in particular identifiable ICE/IME genes often located farther from ISs than prophages, suggests the existence of still poorly understood mechanisms specifying the spatial organization of the ecological niches of ISs and, more generally, of MGEs. The widespread presence of terminator-like sequences upstream of IS CDSs, which are found both within the IS itself and just outside it, also suggests the existence of regulatory mechanisms mitigating IS transposase expression. This regulation could prevent excessive transposition activity, which might otherwise lead to an explosion of IS copies detrimental to the host genome (3), and, hence, contribute to the survival of ISs within genomes. It is important to emphasize that the observed genomic patterns ultimately result from selective processes. This is particularly evident in the fundamental genomic properties associated with IS niches. Specifically, the observed bias of IS niches towards intergenic regions and areas that do not disrupt the functional organization of genomes supports a major role of purifying selection in shaping IS distributions. Nevertheless, our finding that ISs more frequently disrupt genes involved in defense mechanisms, DNA repair and extracellular structures also suggests that some events may involve positive selection. It is also possible that these contexts favor the survival of the ISs themselves.

The specific depletion of GC content in the vicinity of ISs – strongest near the IS and extending up to 100 kb – introduces a new dimension to the discussion on the origins of the heterogeneity of the GC content across bacterial genomes (43) and within them. Namely, a commonly proposed explanation for intra-genome heterogeneity is that genes with higher recombination rates tend to have higher GC content (44; 45), and that recombination is often more frequent in core genes than in accessory genes (see e.g. (46; 47; 48)). Yet, the underlying reason why recombination is GC-biased remains actively debated – see e.g. (49). Some have argued for a mechanistic bias inherent to the recombination process itself (44), whereas others have argued for a selective pressure favoring optimal expression of conserved genes (50; 45; 49; 51). Here, we first note that HGT-associated hotspots have been shown to harbor higher recombination rates (12). Under a purely mechanistic hypothesis, one would thus expect an enrichment of GC content, at least in the intergenic regions near ISs, given that many of these ISs are found within such hotspots (12). However, we observe the opposite: a pronounced GC depletion, for both non-coding flanking regions and CDSs, and irrespective of codon position for the latter. This contradiction therefore supports the alternative explanation of a relaxed selective pressure within IS-associated niches. Yet, it remains unclear how this could also account for the AT enrichment specifically observed in intergenic regions. One possibility is that part of the signal arises from an enrichment of AT-rich subsequences that are prone to gene silencing and involved in the regulation of horizontally acquired genes (52), given that xenogeneic silencing has been shown to both act as a selective force shaping the GC content of genomes (53) and to influence the ecological niches of ISs (24).

Altogether, our findings point to a situation involving both physical and ecological isolation of ISs, and more broadly of MGEs. Physical isolation refers to the tendency of these elements to localize in specific regions of the host genome, whereas ecological isolation, also called isolation by environment (54; 55), would reflect selective pressures in these regions that differ from those acting on the rest of the genome.

Beyond the niche characteristics shared by all IS families, our results also reveal that some families exhibit distinct niche specificities with respect to flanking gene synteny, distance to neighboring genes, or the upstream presence of transcription terminators. Because some of these specificities are expected to depend on the tool used to identify ISs, we have verified that the main findings we report remain robust when switching between annotation tools (see Supplementary Note 3). In this context, IS-specific properties are likely linked to family-specific target sequence preferences, as well as to specific expression properties and functional roles. This is further supported by the consistency between some of the patterns identified in our large-scale study and behaviors reported in more specific studies. For instance, members of the IS*30* family – one of the few families that insert between syntenic gene pairs more frequently than expected – have been reported to activate neighboring genes by creating hybrid promoters (10), which might ultimately confer a benefit to the host. Another example is the IS*481* family, in which many elements are found juxtaposed to flanking CDSs: a similar bias was recently reported in *Bordetella pertussis* and shown to be associated with the regulation of gene expression in the flanked gene (32). Finally, our identification of terminator-like sequences both downstream of non-IS CDSs and upstream of specific IS families strongly suggests that these sequences indeed function as transcription terminators potentially attenuating the expression of the IS transposase, as demonstrated in a previous work on the IS*200*/IS*605* family (33). Demonstrating the functional impact of all identified patterns in a systematic manner may nevertheless prove challenging, as it would require both a refined case-by-case analysis of genomic sequences and experimental testing of the resulting predictions.

From a broader perspective, by framing IS biology within a quantitative ecology of transposons, we anticipate new lines of investigation into their evolutionary dynamics, with new types of questions such as: To what extent do ISs compete with one another? Can ISs specifically collaborate, either with each other or with other MGEs? Is there a process akin to ecological succession when IS elements and other MGEs colonize new genomic regions? Pursuing these directions will not only clarify the roles of ISs and MGEs in genome evolution but also help establish a unifying ecological framework for understanding the mobile genome.

## Materials and Methods

### Genome data and statistics

30,499 full prokaryotic genomes were downloaded from NCBI on July 15, 2022. We used the taxonomy and replicon classification (chromosomes/plasmid) provided in the associated metadata. The dataset includes 8,106 distinct species. We accounted for sampling bias by weighting each genome by the abundance of the species (weight = 1/(nb of genomes)) and by conducting analyses using these weights. Genomes lacking species taxid information were dropped from the analysis (90 genomes). In specific analyses where weighting is cumbersome, we consider a single strain per species, as in Fig. 1.

### Detection of ISs

ISs were detected using the digIS software (https://github.com/janka2012/digIS) (23), which requires only the fasta files of genomes – a cross-validation was performed by comparing digIS outputs with those of ISEScan (22) for a subset of 21,298 genomes (see Supplementary Note 3). From a total of 1,502,064 reported potential ISs, we retained 993,902 ISs reported as complete elements. Among these, 770,751 elements were found on chromosomes, 153,409 on plasmids, and 69,742 on unclassified replicons. These elements fall into 30 families, corresponding to the ISfinder classification (56). Some detected IS elements matched several HMM models: when the models belonged to the same family, they were classified into it; otherwise, they were labeled as “other” and excluded from our analysis of family-dependent properties.

### Rank abundance curve (RAC) and ecological models

We considered two generic ecological models – lognormal and power law – to fit the RAC. The expected abundance *f* (*r*) for a given rank *r* in these models is given by: 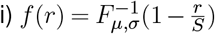 for the lognormal model, where *F*_*µ,σ*_ is the cumulative density function of the lognormal distribution with mean *µ* and standard deviation *σ, S* is an effective maximum number of species, and ii) *f* (*r*) = *Ar*^*−α*^ for the power-law model. The model parameters (*µ, σ*, and *S* for the lognormal model; *A* and *α* for the power-law model) were estimated by least-squares minimization.

### Identification of insertion context

The 200 bp flanking ISs on each side were extracted and joined into a single sequence, which was subsequently searched against the NCBI prokaryotic sequence database using MegaBLAST (with an e-value threshold set at 1e-60 and `max_target_seqs` at 5). When a hit sequence was available in our database, the annotation file for that sequence was examined to see if the hit position was annotated as a coding sequence (CDS). If so, the IS was classified as having interrupted a gene. Otherwise, it was classified as an intergenic transposition.

### Identification and analysis of genes flanking intergenic ISs

For each detected IS (except ISs classified as intragenic), we retrieved the flanking genes as the closest upstream and downstream genes using NCBI metadata annotation tables. To avoid issues related to potentially incomplete genome annotations, only ISs with flanking genes located less than 1 kb away were considered for the analysis of gene orientation and synteny (640,578 ISs out of 993,902).

### Functional annotation of genes using COG categories

The functional classification of interrupted and flanking genes was performed using COGclassifier (https://github.com/moshi4/COGclassifier), which classifies genes based on sequence similarity to representative orthologs. We retained 22 of the 26 COG categories (57), excluding categories A, B, Y, and Z (eukaryote-specific).

### Operon database and gene synteny

*E. coli* K12 MG1655 operons were retrieved from RegulonDB v12.0 (58). Genes in a synteny relationship were identified on the basis of their COG annotation (see above), using a previously established list of COG pairs identified to be in synteny relationships (59). This list is provided in Supp. Table 2.

For the synteny analysis of gene pairs from distinct operons flanking an IS, we considered only those pairs that occur as neighbors in at least one other *E. coli* genome.

### Identification of terminator-like structures

To identify potential Rho-independent intrinsic terminators around ISs, we scanned upstream and downstream for inverted repeats followed by T-tracks. Based on structural analyses (60; 61), we considered inverted repeats with gap sizes ranging from 3 to 7 (hairpin loop size in bases) and arm lengths from 4 to 14 (stem size in base pairs), and T-tracks composed of more than 4 Ts. The terminator analysis was conducted around complete ISs (as defined by digIS) and around IS CDSs (identified by intersecting digIS coordinates with NCBI CDS annotations). For comparison, we also scanned sequences flanking randomly sampled CDSs from the genomes of our dataset.

### Identification of IS clusters

Chromosomal ISs were considered part of the same cluster if they were adjacent along the DNA and separated by less than 2.5 kb, the typical maximum distance observed to flanking CDSs (Fig. S4A). To mitigate false negatives, when two ISs were not immediate neighbors but were separated by less than 2.5 kb with a transposase annotated in-between them (despite not appearing in our dataset), the two ISs were considered to be part of the same cluster. We also verified that our findings do not depend on this assumption.

### *E. coli* phylogenetic tree

To examine the dynamics of a specific IS cluster in *E. coli*, we focused on one conserved gene pair (as defined by PpanGGOLIN; see below) and randomly sampled 50 genomes containing an IS cluster between these genes, along with 30 genomes devoid of IS at that locus. A phylogenetic tree was then built for these 80 genomes using 41 phylogenetic markers selected with the markerfinder tool (see methodology described in (62)), with a *Xanthomonas oryzae* genome (GCA_001928215.2_ASM192821v2) used as an outgroup to root the tree (removed on the final visualization). The tree was simplified to keep only 59 leaves.

### Persistent, accessory genes and regions of genome plasticity (RGPs)

Persistent, shell, and cloud genes were identified using the PpanGGOLiN tool (36) (https://github.com/labgem/PpanGGOLiN). For each chromosome in the genomes of the three analyzed species (*Escherichia coli* (GC%=50.7), *Pseudomonas aeruginosa* (GC%=66.2), and *Acinetobacter baumannii* (GC%=39.1)), PpanGGOLiN reports segments classified as RGPs (regions of genomic plasticity). Chromosomes were then divided into 1,000 bins, and the probability to be in a RGP across all chromosomes was computed for each bin.

### Profiles of IS occurrences along a normalized chromosome

For the three species *E. coli, P. aeruginosa*, and *A. baumannii*, containing respectively 1,917, 451 and 444 genomes in our dataset, chromosomes were first reoriented so that the origin of replication (*oriC*) corresponds to position 0, with the *rplS* gene always located on the right replichore. Second, for each chromosome, IS positions and other genomic features (RGPs, persistent genes, and so on) were normalized to chromosome length, such that position 0.5 corresponds to the replication terminus (*ter*) and position 1 loops back to *oriC*. To avoid overcounting IS occurrences inherited from a common ancestor, those with identical flanking genes and orientation were weighted so that their total contribution summed to 1.

In this context, the profiles in Fig. 5A, Fig. 6, Fig. S11, Fig. S12, and Fig. S13 were computed by dividing chromosomes into 1,000 bins (approximately 5 kb, 4 kb, and 6.5 kb resolution for *E. coli, A. baumannii*, and *P. aeruginosa*, respectively). For clarity, curves were smoothed using a Gaussian filter with a standard deviation of 2 bins to reduce small-scale fluctuations, as the main variations occur at a much larger (∼100 kb) scale.

### GC content deviation profiles around IS positions

For all genomes in our dataset, the deviation from the average GC content around ISs was computed at 1,000 positions logarithmically spaced from 1 bp to 1 Mb away from each IS, both upstream and downstream the IS with respect to its transposase CDS reading orientation. A corresponding profile was computed around an equal number of randomly selected chromosomal CDSs in each genome. Similarly, we computed the profiles associated with different genomic features (intergenic regions, CDSs, as well as first (GC1), second (GC2), and third (GC3) codon positions) by retaining only positions belonging to the investigated feature. Note that in this case, the average GC content was computed using only the positions associated with the feature. Note also that the analysis considering three groups of genomes (Fig. S16) correspond to three distinct regimes of GC3 levels (GC1 is always higher than GC2): (i) GC3 is the lowest; (ii) GC3 is intermediate (between GC2 and GC1); and (iii) GC3 is the highest.

### Detection of MGEs

Prophage CDSs were detected using the DB-SWA tool (63) (https://github.com/HIT-ImmunologyLab/DBSCAN-SWA/). ICEs and IMEs were detected using the CONJScan (v2.0.1) module of MacSyFinder (v2.1.3) (64; 65) (https://github.com/macsy-models/CONJScan). Integrons were detected using IntegronFinder (66) (https://github.com/gem-pasteur/Integron_Finder). For each of these categories, a mask was applied to the genomes to consider only relevant positions when analyzing the frequency and characteristics of each type of MGE around IS positions. To make sure masks were exclusive, we defined ICE annotation as superseeding integron annotation, integron annotations as superseeding prophage annotations, and prophage annotations as superseeding IS annotations. We also verified that our results do not depend on sequences with ambiguous annotations (*i*.*e*., sequences annotated as at least two different types of MGEs.)

## Acknowledgements

We would like to thank the GeDy team at the CBI in Toulouse for their constructive feedback on the first version of the manuscript and for insightful discussions, particularly Jean-Yves Bouet, Manuel Campos, François Cornet, Catherine Guynet, and Patricia Siguier. IJ also thanks Olivier Espeli and Olivier Rivoire for discussions and comments.

We acknowledge the use of AI (Mistral Emmy, CNRS) to polish the English of the manuscript.

## Data availability

A repository containing the scripts used to generate the results of the article can be found at https://zenodo.org/records/20623700.

The public data used in this study are not shared, as the corresponding compressed files are too large (several hundreds of GB). The information required to access these data is provided in the Methods section of the article.

## Supplementary Information

### Supplementary Note 1: Random location models for the formation of IS clusters

We test whether the clustering properties of ISs can be explained by models in which their genomic locations are randomly assigned under three different scenarios: (I) a model where the entire chromosome is accessible (according to a uniform probability distribution), (II) a model where IS-accessible regions are limited to a fraction of the chromosome, and (III) a model where the number of accessible sites scales linearly with the number of ISs.

### General framework

Let us first consider a chromosome containing *n* ISs plus *A* accessible sites with which ISs can be exchanged, i.e. the permutation of their location does not alter the selective pressure acting on the host. Considering ISs as occupied sites, this means that, for a given configuration of IS locations, there are *n* occupied and *A* empty accessible sites (see panel A of the associated figure). It also means that any permutation of the IS positions among these accessible sites corresponds to a valid configuration that can be sampled during the course of evolution. One can then ask: what is the probability *P*_*n,A*_(*k*) that a given cluster has size *k* (including the case *k* = 1), assuming an equiprobable distribution over all possible configuration of IS locations? *P*_*n,A*_(*k*) can be computed by noting that it corresponds to the probability that an occupied site, flanked on its left by an empty site, is followed on its right by *k* − 1 occupied sites and an empty site at the *k*^th^ position. Defining 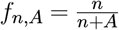 as the fraction of accessible sites that are occupied, we have:

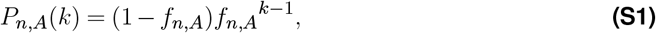

a formula that can be verified by simulating the (evolutionary) sampling of all possible configuration of IS locations (see panel B of the associated figure).

The corresponding mean size of the clusters, *m*_*A*_(*n*), as a function of *n* is given gy:

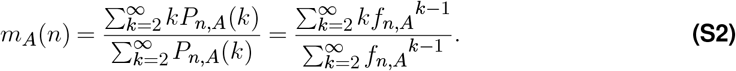

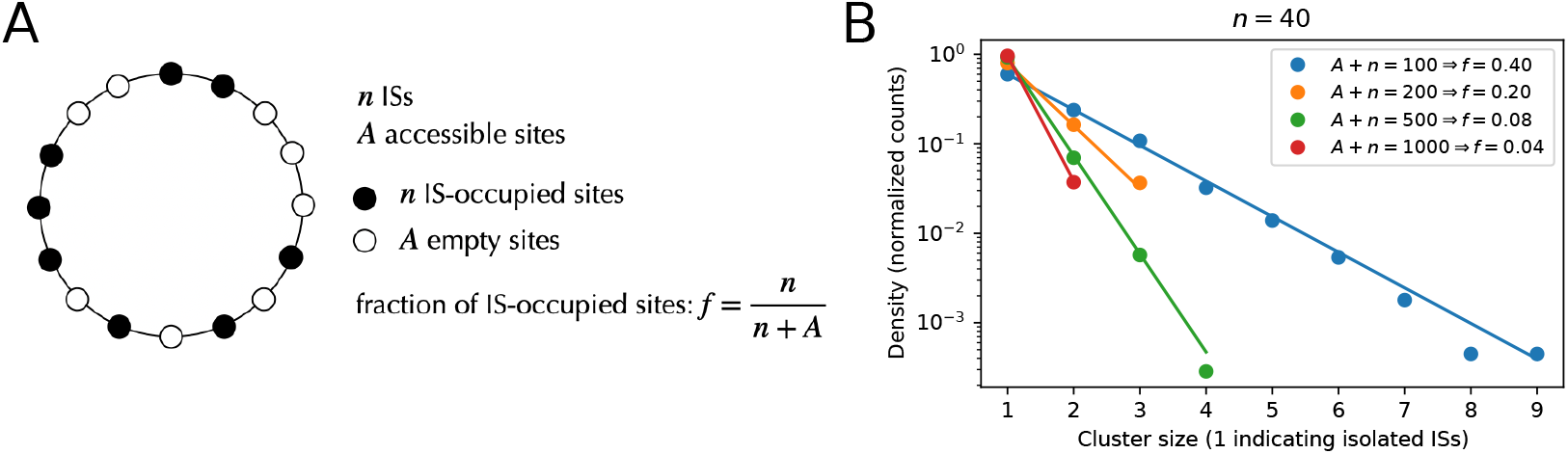

A. **Scheme of the random location models**. B. **Comparison between simulations (points) and analytical predictions (lines) given by Eq. S1**.

Dropping the subindexes of *f*_*n,A*_ for lightness, we obtain:

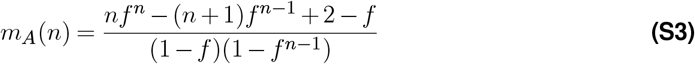

In particular, for values of *n* such that *nf*^*n−*2^ ≪1, which occurs typically at *n* ≃10 in the case of the fit to the genomic data (see Fig. 4C), one has:

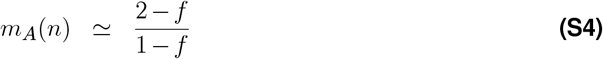

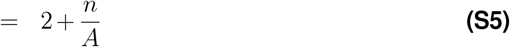

#### Scenario I: full access to the chromosome

In this scenario, all ISs can be located at any position along the chromosome. In the terminology of our above general model, this means that IS locations can be exchanged with any other gene, such that *n* + *A* represents the chromosome length (in number of genes). For simplicity, in Fig. 4C, we used *A* = 4000, which corresponds to the typical length of the chromosomes analyzed. Nevertheless, one can compute the mean cluster size *m*(*n*) as a function of *n* by considering the specific length of each chromosome. In this case, calling *N*_*C*_ the total number of genes (excluding ISs) and *n*_*C*_ the number of ISs in a chromosome *C*, one has:

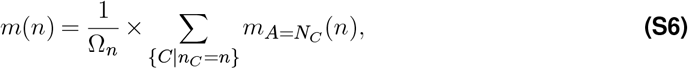

where {*C*|*n*_*C*_ = *n*} denotes the set of chromosomes *C* for which *n*_*C*_ = *n*, Ω_*n*_ = |{*C*| *n*_*C*_ = *n*}| is the number of such chromosomes, and 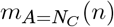 (*n*) is given by Eq. S3. Plots then show that the two approaches yield very similar results (blue curves in the associated figure).

#### Scenario II: limited access to the chromosome

In this scenario, only some part of the genome can be accessed by ISs. In the terminology of our general model, this means that A represents some part of the genome. One can then consider two distinct situations: i) *A* is a fixed quantity, independent of the genome length; ii) *A* is a fraction of the total genome. For simplicity, in Fig. 4C, we used a fixed quantity *A* = 5.4 × 120 as the maximum size of the accessible genomes coming from scenario III (see below). Nevertheless, part of the large variation of data for *n* ≥ 120 can be explained considering Eq. S6 with *A* that is a fixed part of the genome, i.e. using *A* = *αN*_*C*_ such that:

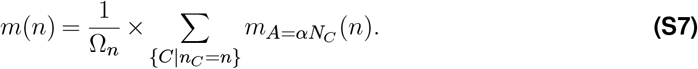

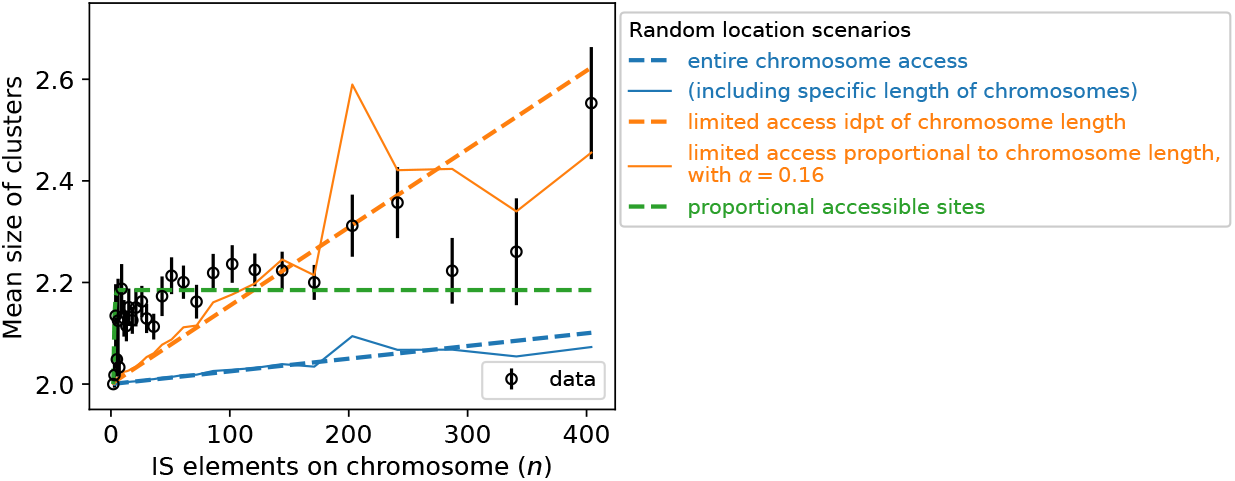

**Effect of considering the specific length of each chromosome in models (solid curves) compared to considering a single average value (dashed curves)**. The model of proportional accessible sites is not concerned since the parameter of the model is independent of the length of the chromosome.

In particular, using *α* = 0.16 provides a reasonable fit of the data (orange curve in the associated figure).

#### Scenario III: the size of IS niche scales linearly with the number of ISs

In this scenario, the additional presence of an IS is accompanied by new genes with which ISs can be exchanged. In the simplest case, we can assume that the number of new accessible sites is proportional to the number of additional ISs such that the size of the IS niche scales with the number of ISs, i.e., *A* = *βn* with *β* ≥ 0. In this case, one has:

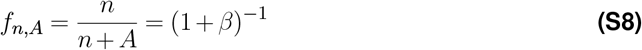

which is independent of both *n* and the size of genomes. Note, then, that in this case, the size distribution of clusters follows an exponential distribution that is independent of these factors as well, just a observed in experimental data (see Fig. 4B).

Regarding the mean size of clusters, we obtain:

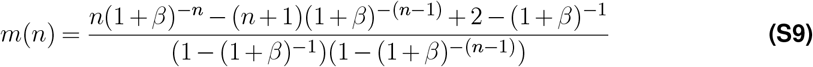

with, in the limit of large *n*:

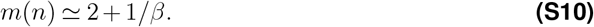

This scenario, with *β* ≃ 5.4, explains particularly well the data for *n*≤ 120 (green dashed curve in the associated figure; see also Fig. 4C).

### Supplementary Note 2: Quantifying the contribution of features to the GC depletion signal

To quantitatively assess the contribution of CDS and intergenic regions to the GC depletion signal, we generated GC content profiles separately for CDS and intergenic positions, using NCBI metadata available for each genome. In this context, as demonstrated below, the contribution of a feature-specific signal at position *x* reads:

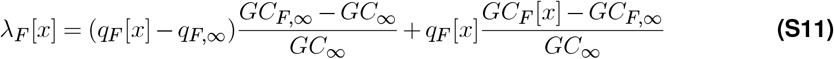

where, for a given feature *F* (either CDS or intergenic): *GC*_*F*_ [*x*] is the corresponding GC content profile, *GC*_*F,∞*_ is the corresponding average GC content at positions far from the IS (in practice, at a distance of 1 Mb), *q*_*F*_ [*x*] is the fraction of times *F* is found at position *x*, and *q*_*F,∞*_ is the fraction of times *F* is found at positions far from the IS. *GC*_*∞*_ is the overall, i.e. feature-independent, average GC content at positions far from the IS.

This contribution consists of two terms that can be analyzed separately. The first term, which equals 0 if *q*_*F*_ [*x*] = *q*_*F*,_ *∞*, reflects either an under-representation (*q*_*F*_ [*x*] *< q*_*F*,_ *∞*, as observed for CDS) or an over-representation (*q*_*F*_ [*x*] *> q*_*F*,_ *∞*, as observed for intergenic regions) of the feature *F*. The second term, which equals 0 if *GC*_*F*_ [*x*] = *GC*_*F*,_, reflects an AT-enrichment if *GC*_*F*_ [*x*] *< GC*_*F*,_ *∞*, which is observed for both CDSs and intergenic regions. In the panels B and C of Fig. S15, we therefore computed the relative contribution of each of these terms to *λ*_CDS_[*x*] and *λ*_intergenic_[*x*], respectively, up to 10 kb, where the GC depletion signal remains clearly visible (see Fig. 7A).

#### Application to the depletion signal around ISs

For Fig. S15A, we applied Eq. S11 using the set of features =ℱ {intergenic, CDS}. Note that for both intergenic and CDS features, the two terms in Eq. S11 are negative, allowing to assess the relative contribution of each term, as done in Fig. S15BC. It is then interesting to note that the first term is negative for different reasons: CDSs tend to be both under-represented (*q*_CDS_[*x*] *< q*_CDS,*∞*_) and more GC-rich (*GC*_CDS,*∞*_ *> GC*_*∞*_), whereas intergenic regions tend to be over-represented (*q*_intergenic_[*x*] *> q*_intergenic,*∞*_) and more AT-rich (*GC*_intergenic,*∞*_ *< GC*_*∞*_). By contrast, the second term is negative for both CDSs and intergenic regions due to their AT-enrichment, i.e., *GC*_*F*_ [*x*] *< GC*_*F,∞*_ for *F* ∈ {intergenic, CDS}.

For Fig. 7D, we quantify the contribution of known MGEs by applying Eq. S11 in the context of CDSs only, separating the CDS of MGEs from the rest. The set of features in this context is therefore ℱ = {CDS(IS), CDS(phage), CDS(ICE/IME), CDS(integron), CDS(other)}.

#### Demonstration of Eq. S11

We consider a complete set of disjoint features ℱ such that, denoting by *q*_*F*_ [*x*] the probability of finding a feature *F* at a genomic position *x*, we have:

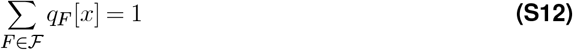

Now, what is the GC content at position *x*? Or, equivalently, what is the probability, denoted hereafter *GC*[*x*], of finding a GC base pair? By definition, we have:

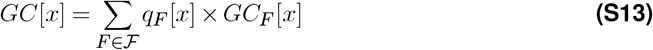

where *GC*_*F*_ [*x*] denotes the GC content associated with feature *F*, or equivalently, the (conditional) probability of finding a GC base pair given that feature *F* is present.

Next, let• *∞* indicate the value of the quantity •when *x*→ ∞. *r*_*GC*_[*x*], the relative GC content at position *x* with respect to the signal at long distances, is defined as:

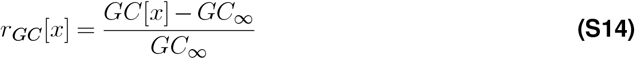

which can also be written as:

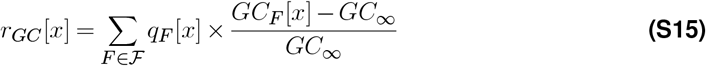

Note here that *GC*_*∞*_ is not necessarily equal to the average value of the GC content over the genome. Actually, it is not in the most general case.

Now, using the fact that *r*_*GC,∞*_ = 0, Eq. S15 leads to:

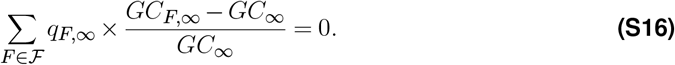

Substracting this equation from Eq. S15, we obtain:

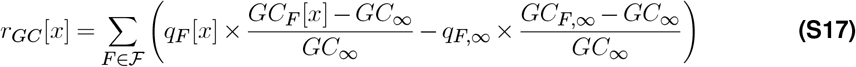

which can be written as:

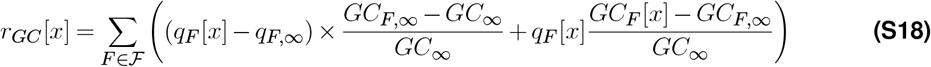

As a result, the contribution *λ*_*F*_ [*x*] of the feature *F* to the relative GC content at position *x* reads:

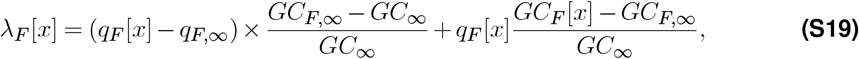

which is Eq. S11.

### Supplementary Note 3: Caveats and limitations

#### Caveats in the analysis of local properties of IS ecological niches

When analyzing the local properties of the ecological niches occupied by ISs within host genomes, it should be kept in mind that, although ISs are typically regarded as elements of recent origin (see e.g. (67)), the observed local context of an IS may not be identical to the ancestral context of insertion, as this context could have changed by subsequent genomic alterations such as deletions, insertions, or rearrangements. Reconstructing the evolutionary trajectories of both ISs and their context is notoriously challenging, if not impossible, and lies beyond the scope of our study. Instead, and despite potential divergence from ancestral states, here we interpret the observed local contexts in which ISs are found as ecological niches in which they are able to survive.

### Cross-validation of digIS outputs

A cross-validation was performed by comparing digIS outputs with those of ISEScan (22) for a subset of 21,298 genomes. Overall, detection of IS elements is consistent between the two tools: the number IS copies detected for each family in each genome is highly correlated (see Figure below). In cases where the tools report widely different numbers of copies, ISEScan tends to report more copies than digIS. For IS*1182* and IS*607* families, the two tools report significantly different results (correlation coefficients of only 0.60 and 0.17, respectively). The manual examination of copies reported exclusively by one tool showed that while IS*1182* copies reported by ISEScan only were presumably true positives missed by digIS, IS*607* copies flagged by ISEScan were most likely false positives (no significant similarity with any copy listed on the ISFinder database (56)). Additionally, digIS also detects ISs belonging to the IS*H6*, IS*Lre2*, and Tn3 families, which is not the case for ISEScan. Because of these findings, we chose to work on digIS outputs: despite the fact that our analysis likely excludes true positives for families other than IS*607*, the significant number of hits reported by digIS was more than sufficient to perform statistically discriminating analyses while avoiding the inclusion of potential false positives.

We also note that some IS-specific properties were not always robust when analyzed ISEScan instead of digIS. In particular, specific peaks in the distances separating ISs from their flanking genes did not always coincide between the two tools. For these properties, the quality of annotation of ISs, as well as the annotation of their limits, plays a crucial role. In the manuscript, we therefore discuss properties and patterns that are observed using either tool.

#### Limitations and robustness of automated detection approaches

The digIS approach, based on HMMER models, offers greater flexibility than BLAST-based methods and is specifically designed to detect distant IS elements. However, it still relies on known, curated IS sequences from ISfinder, introducing a potential bias towards detecting elements similar to those already characterized. Due to the large size of our dataset, manual curation was not feasible, and despite restricting our analysis to complete IS elements, some false positives may remain. Similarly, the identification of MGEs other than ISs (prophages, ICEs and IMEs, and integrons) was automated using DB-Scan SWA, CONJscan, and IntegronFinder, respectively, without manual verification. As a result, the reported MGE distributions within individual genomes may not be entirely precise.

Nonetheless, our goal is not to analyze specific genomes in detail, but to provide a broad overview of IS distribution patterns within and across genomes. Minor inaccuracies in IS and MGE detection are unlikely to significantly influence our findings, a conclusion supported by the consistency of our results with previous large-scale studies of IS distribution. In other words, the trends we observe are robust and unlikely to be driven by isolated detection errors.

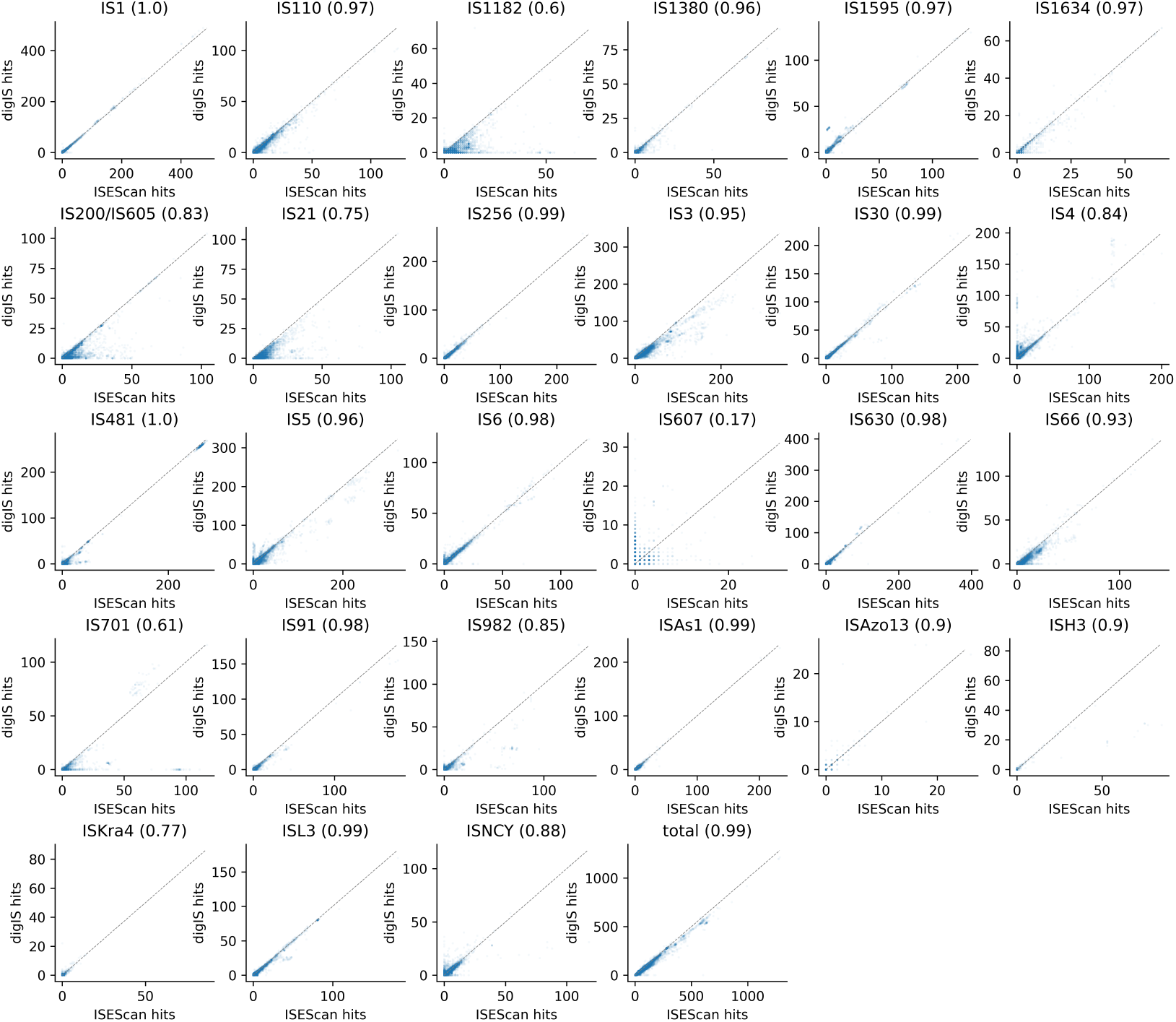

**Cross-validation of digIS hits**. ISEScan and digIS outputs were compared for a subset of 21,298 genomes. Plots show the number of copies detected in a given genome for each family by either ISEScan (x-axis) or digIS(y-axis). Pearson correlations between the two distributions are indicated between parentheses.

## Supplementary Note 4: Supplementary figures

**Figure S1.**
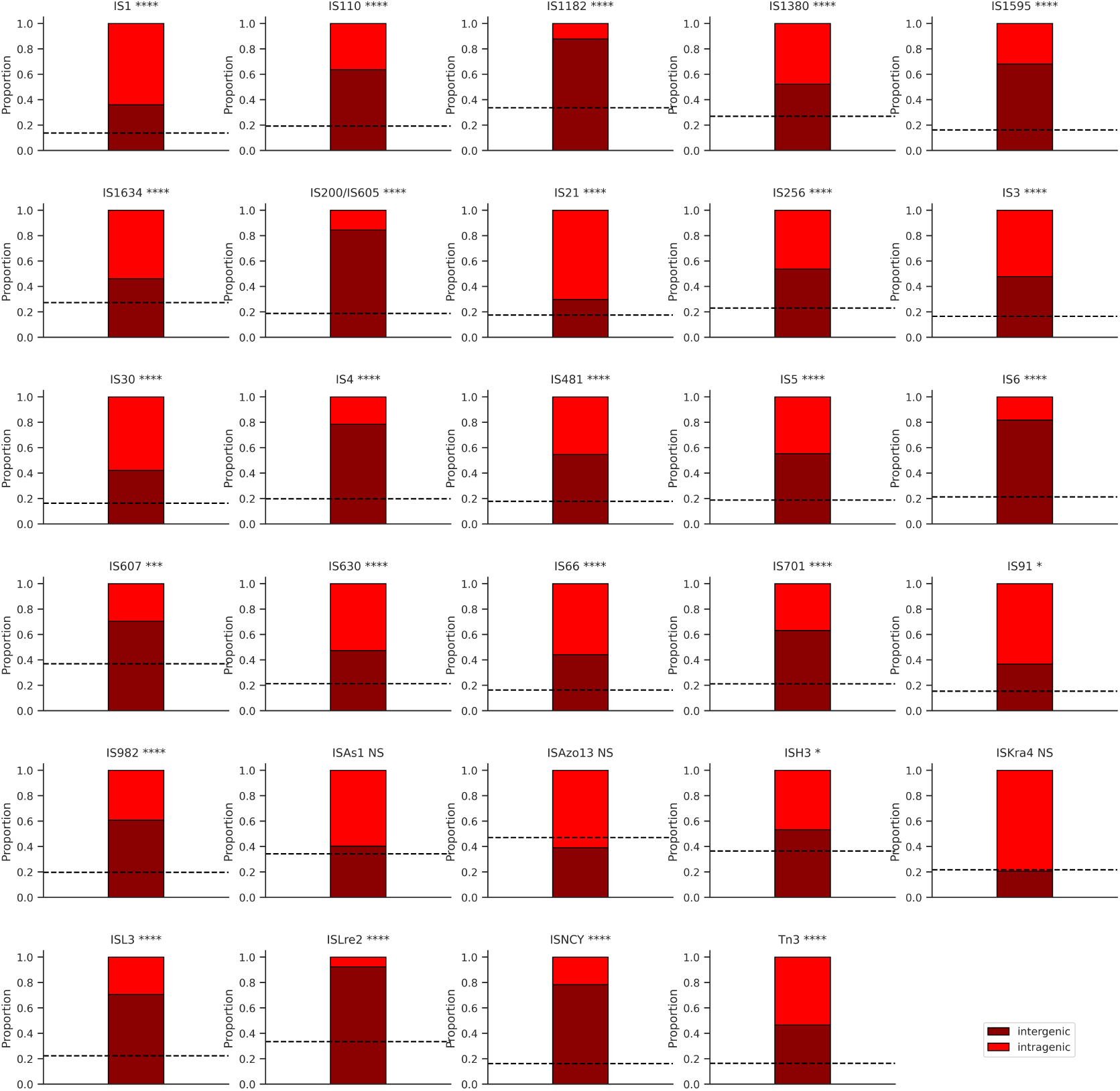
Complementary to Fig. 2A. Proportion of intergenic ISs for each IS family. The whole chromosomal proportion indicated by the horizontal dashed line is computed based on the genomes in which the corresponding IS is present. The ISH6 family is not shown as no data was available. Statistical significance was computed using a Chi2 test. NS: *P* value > 0.05. *: *P* value < 0.05. **: *P* value < 0.01. ***: *P* value < 0.001. ****: *P* value < 0.0001.

**Figure S2.**
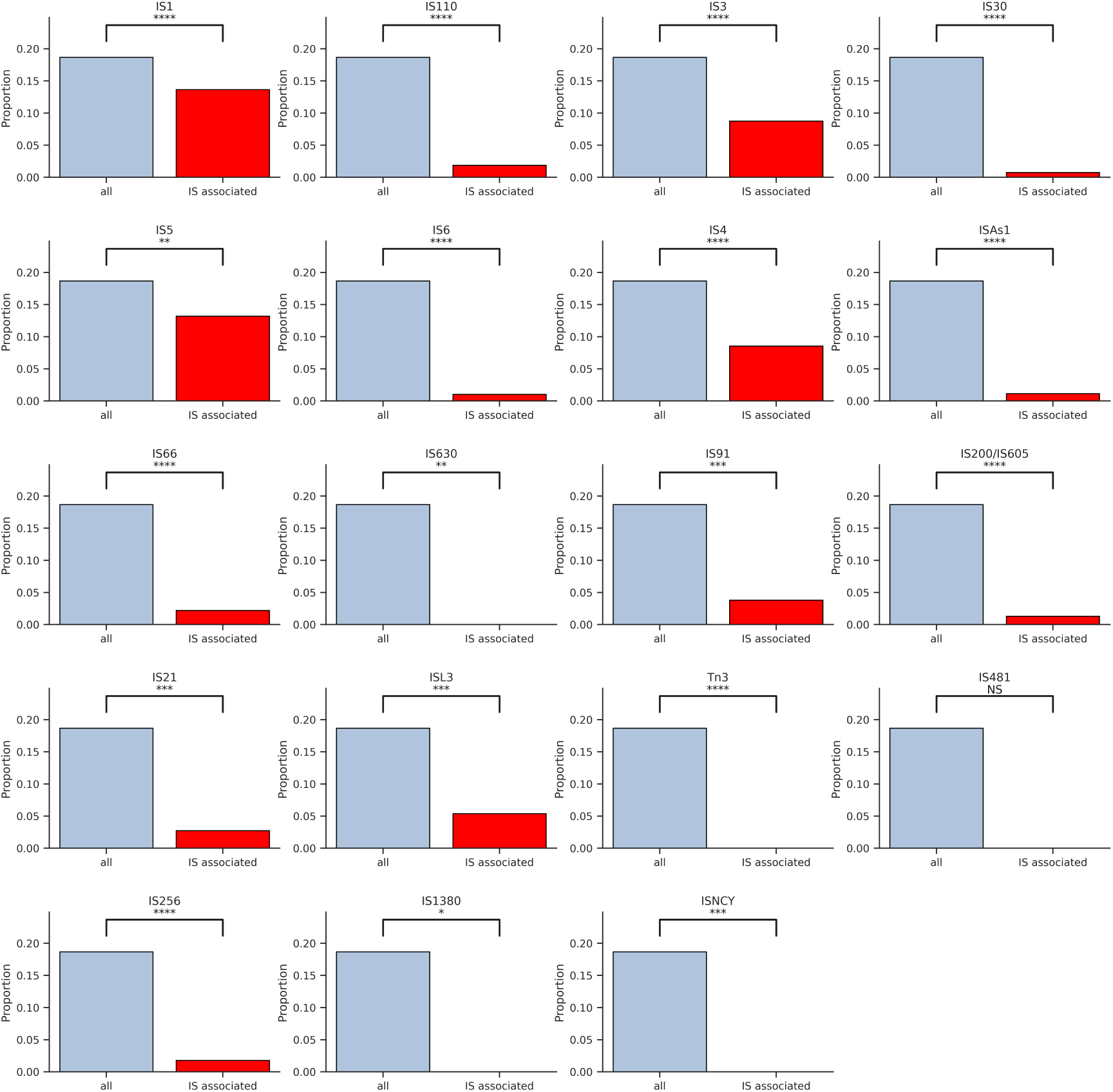
Complementary to Fig. 2B. In *E. coli*, fraction of intergenic regions found inside of an operon (blue) versus fraction of intergenic IS found inside of an operon (red), shown by IS family. Statistical significance was computed using a Chi2 test. NS: *P* value > 0.05. *: *P* value < 0.05. **: *P* value < 0.01. ***: *P* value < 0.001. ****: *P* value < 0.0001.

**Figure S3.**
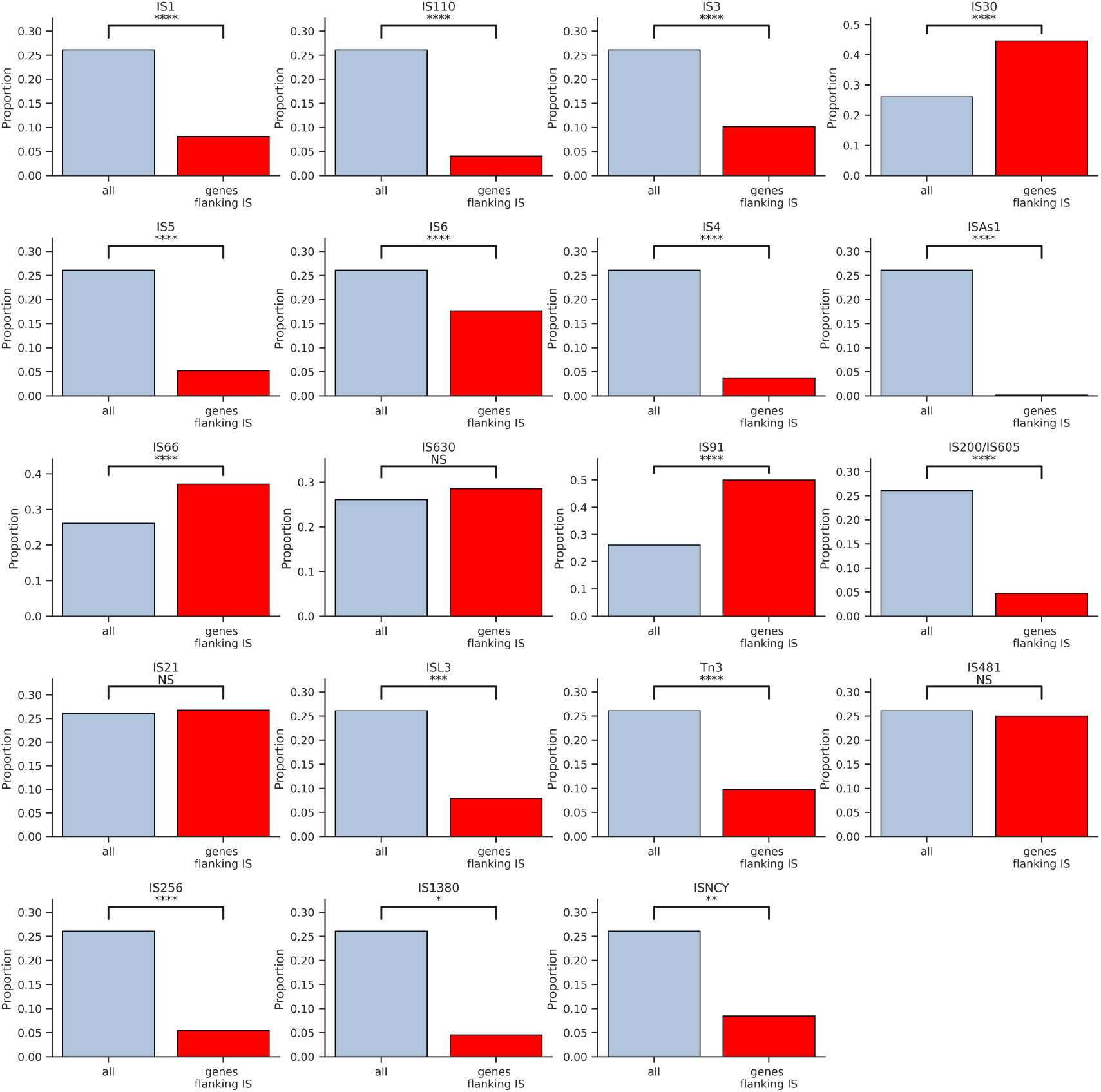
Complementary to Fig. 2C. Fraction of *E. coli* syntenic neighbor genes among genes that do not belong to the same operon in the genome (blue) vs around ISs (red), shown by IS family. Statistical significance was computed using a Chi2 test. NS: *P* value > 0.05. *: *P* value < 0.05. **: *P* value < 0.01. ***: *P* value < 0.001. ****: *P* value < 0.0001.

**Figure S4.**
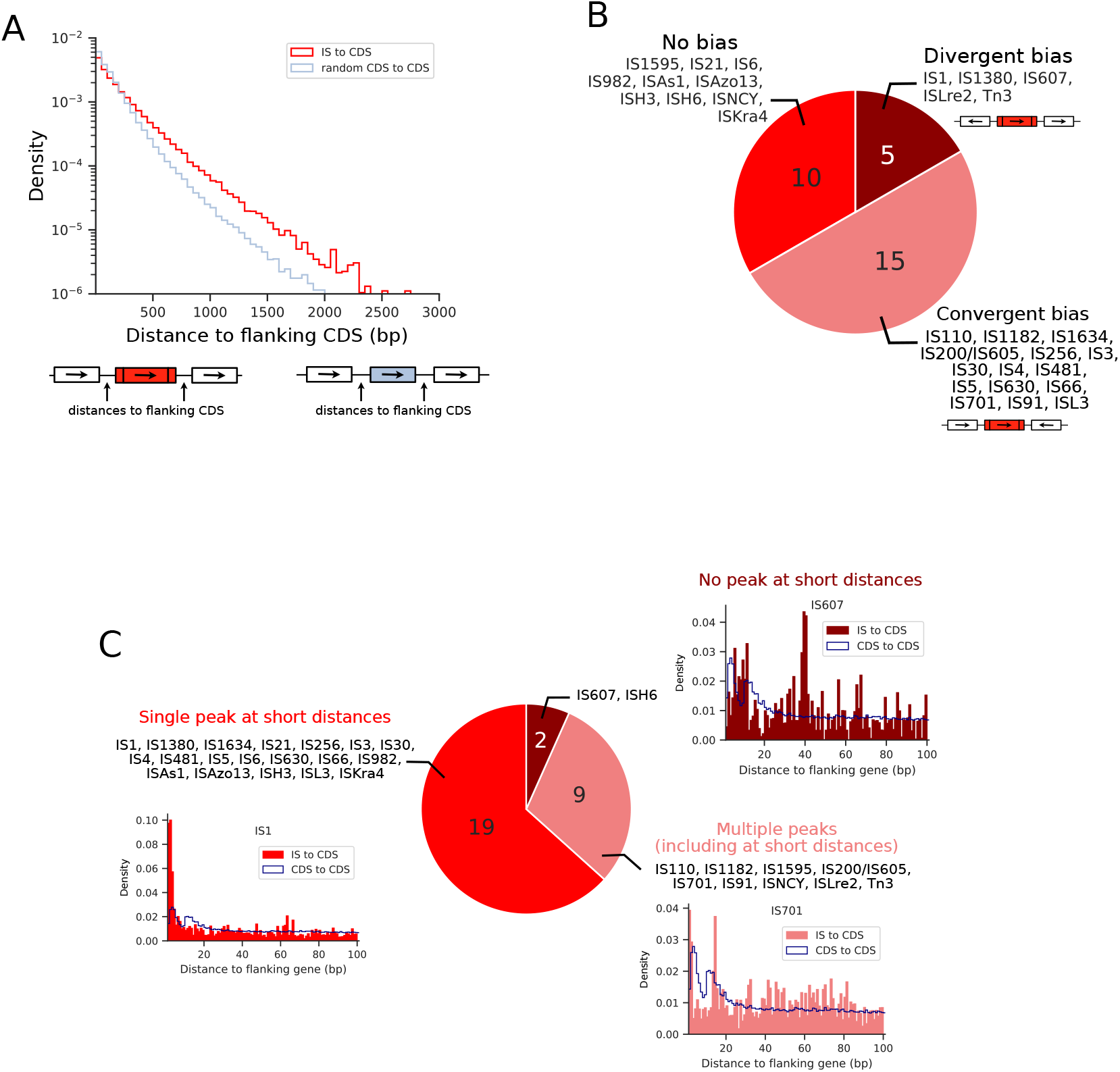
Complementary to Fig. 2. Distance and orientation properties of IS flanking genes. A. Normalized distribution of distances to CDSs flanking ISs (red) or random CDSs (light blue). An analysis of IS flanking regions in genomes lacking the IS further suggests that the greater distances observed with ISs stems from original insertions into longer intergenic regions rather than from post-insertion elongation (Fig. S20). B. Distribution of IS families showing no bias in flanking gene orientation (red), a bias toward divergent flanking genes (dark red), or a bias toward convergent flanking genes (pink). C. The curves show distributions of distances between ISs and the flanking CDSs (red) or between random pairs of neighbor CDSs (blue). The pie chart indicates the IS families showing a single peak at short distances (1–5 bp) (red), multiple peaks including at short distances (pink), or no peak at short distances (dark red), with an example distribution for each category.

**Figure S5.**
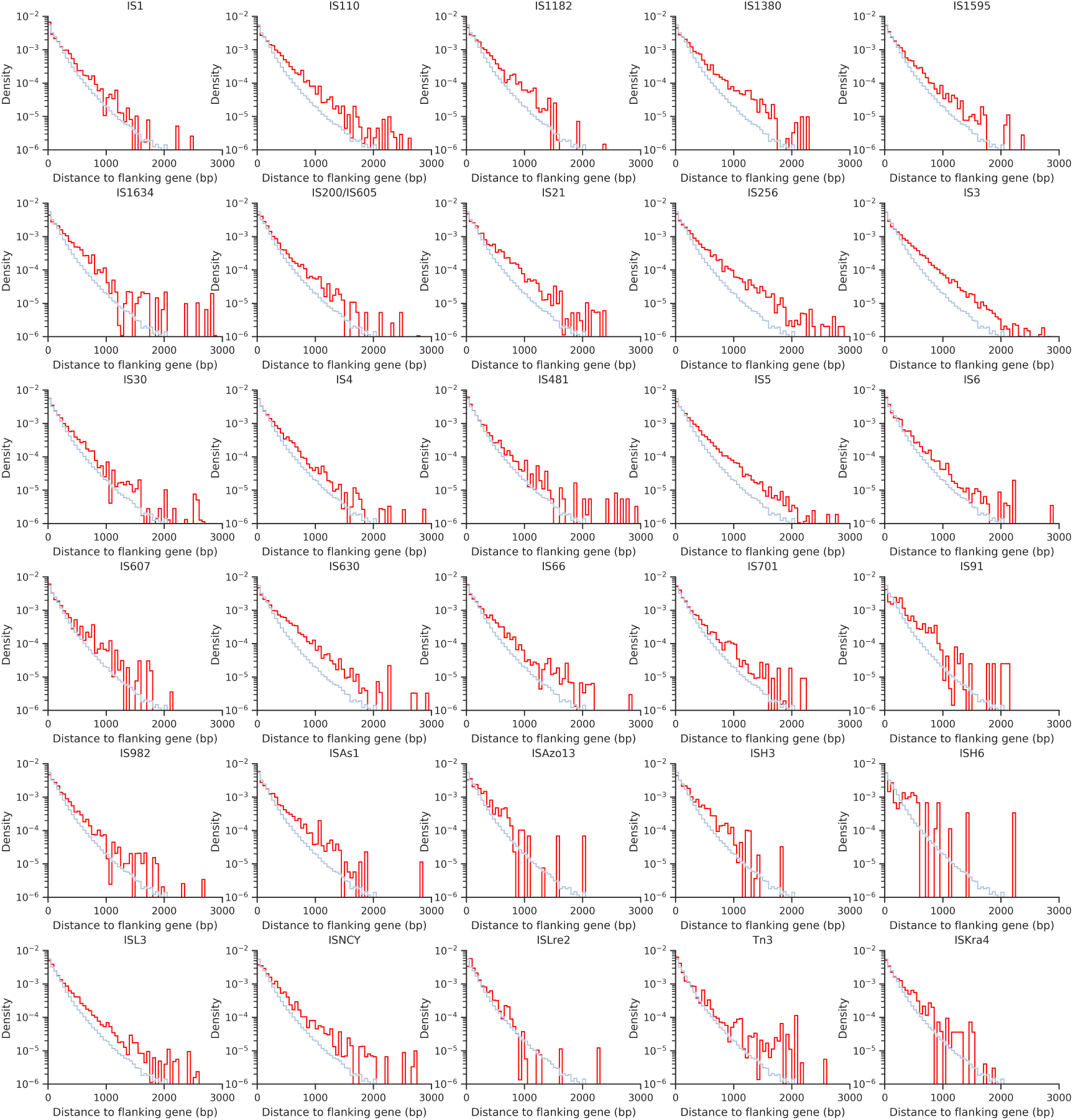
Complementary to Fig. S4A. Normalized histogram of distances to flanking genes for random CDSs (blue) and ISs (red), shown by IS family.

**Figure S6.**
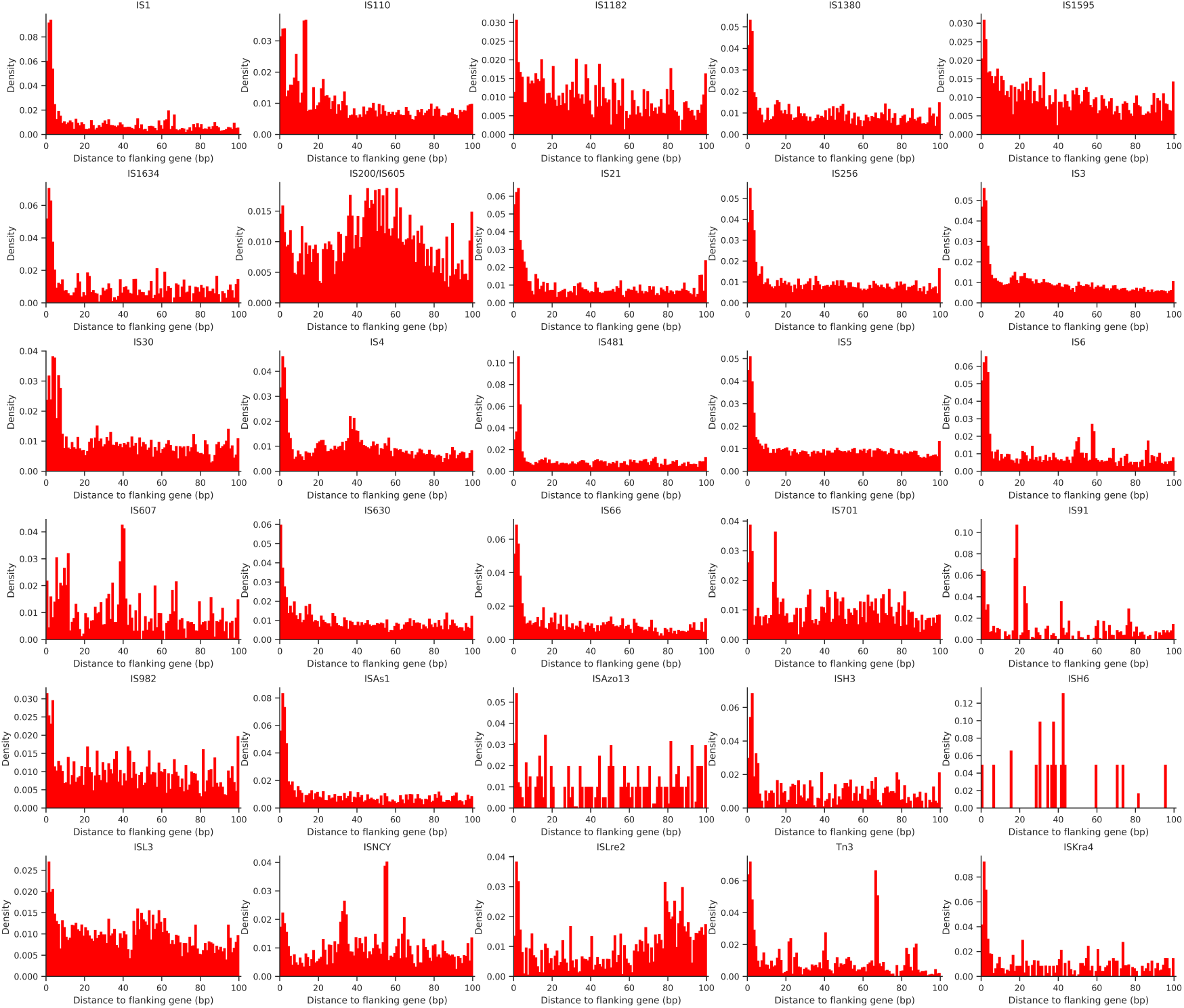
Complementary to Fig. 2E. Normalized histogram of distances below 100 bp to flanking genes of ISs, shown by IS family.

**Figure S7.**
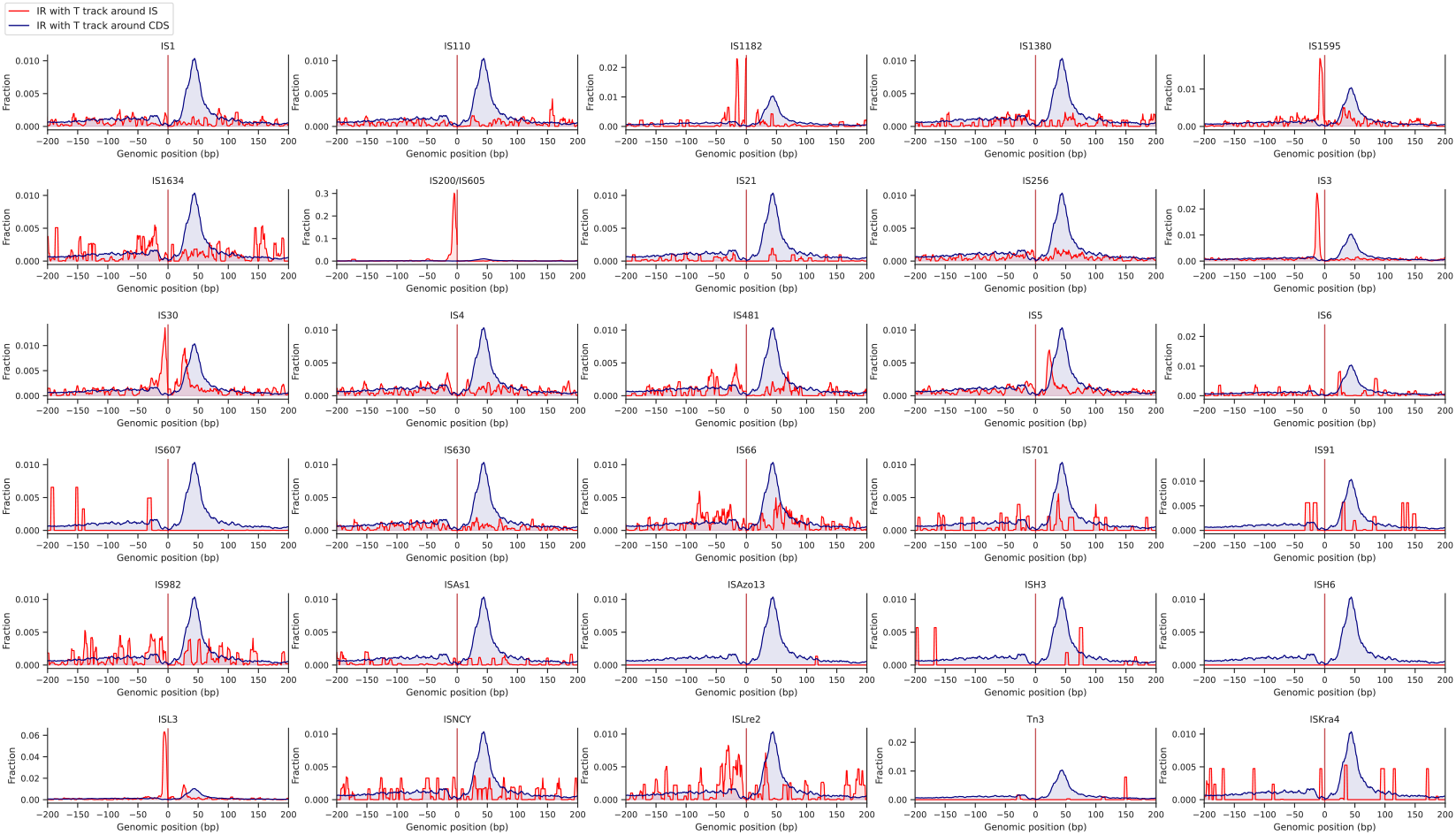
Complementary to Fig. 3. Frequency of terminator-like sequences in the 200 bp upstream and downstream of IS (red) and gene CDSs (blue) shown by IS family.

**Figure S8.**
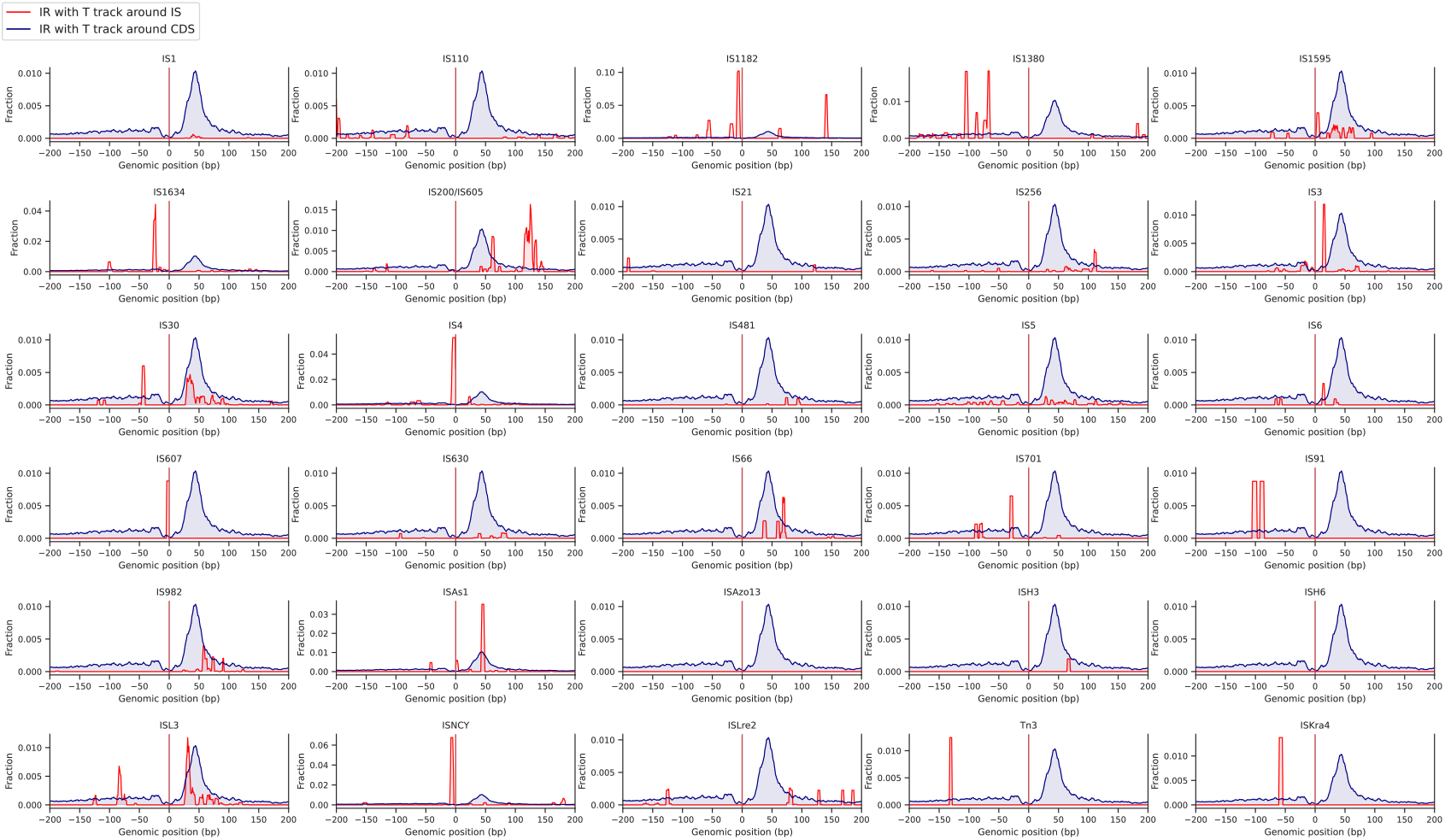
Complementary to Fig. 3. Frequency of terminator-like sequences found inside of IS sequences (between the limits of the IS and the CDS) (red) and around gene CDSs (blue) shown by IS family.

**Figure S9.**
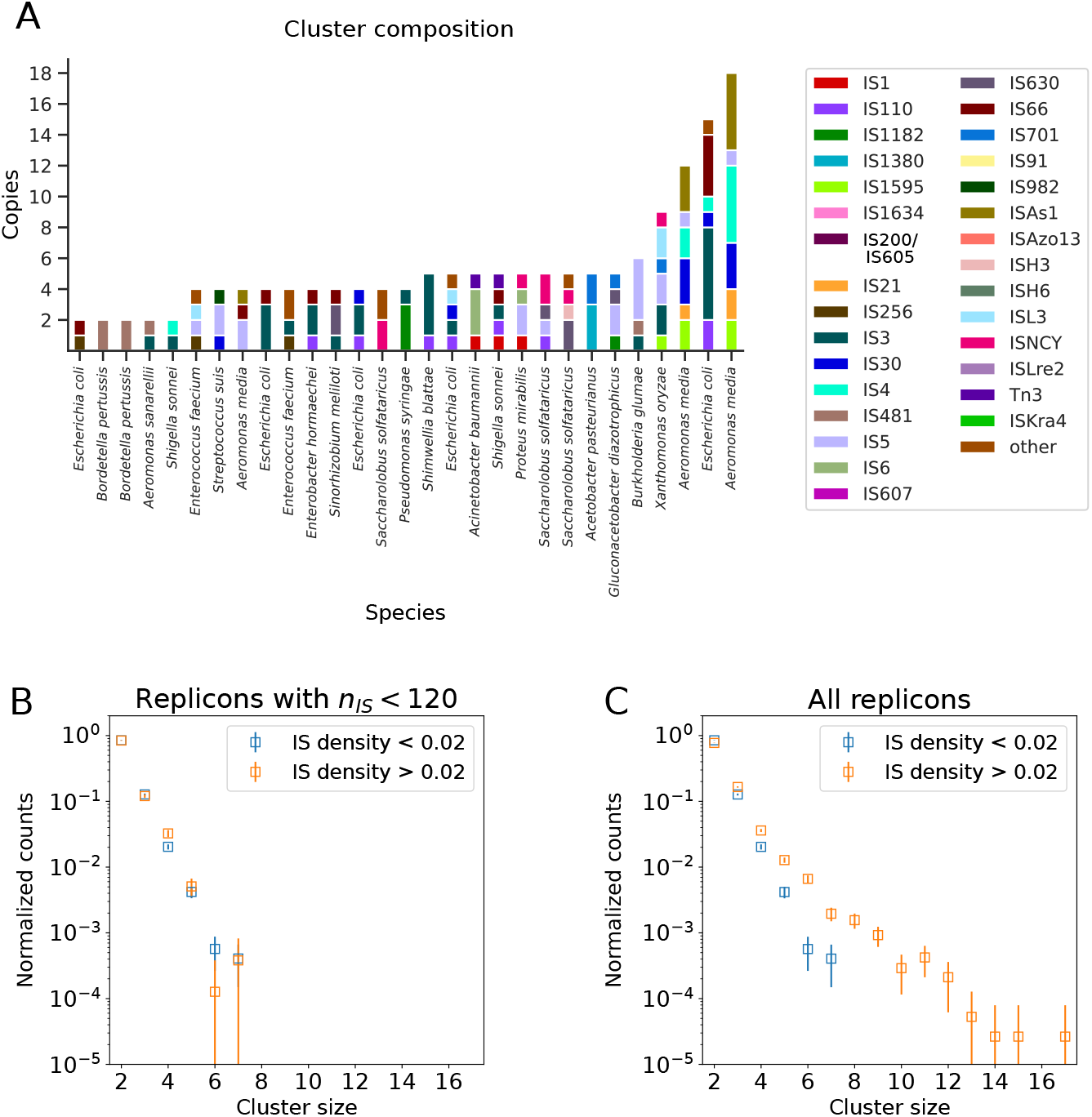
Complementary to Fig. 4. IS clusters: diversity and statistical properties. A. Example of cluster compositions for clusters of different sizes coming from different organisms. B,C. IS cluster size distributions for B) replicons containing fewer than 120 ISs and C) all replicons. The data are shown by separating replicons according to their IS density, defined as the number of ISs divided by the number of genes in the replicon.

**Figure S10.**
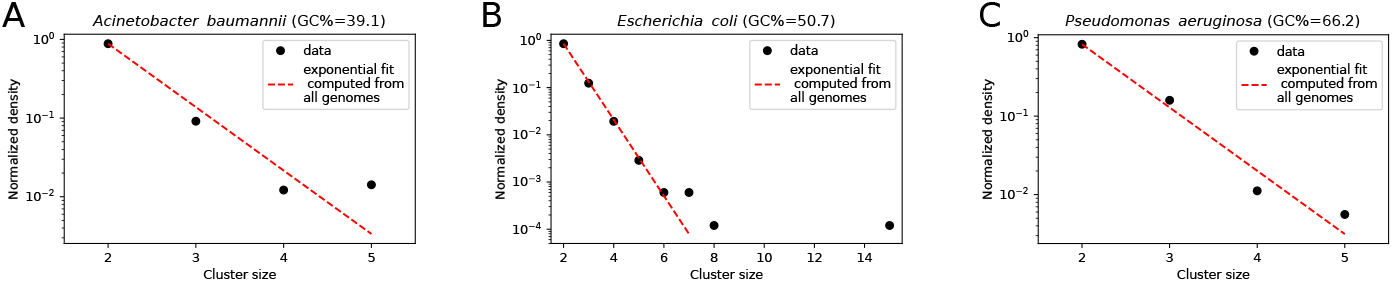
Complementary to Fig. 4. IS cluster size distributions computed over the genomes of three species. Left) *A. baumannii*, GC%=39.1. Center) *E. coli*, GC%=50.7. Right) *P. aeruginosa*, GC%=66.2. The red dashed lines represent the exponential fit obtained by considering the entire set of genomes in our datasets (8,106 species). The apparent differences of slope are due to different ranges of the x-axis. The different x-axis ranges mainly reflect the larger number of clusters in *E. coli*(due to more chromosomes), along with one outlier containing 15 ISs.

**Figure S11.**
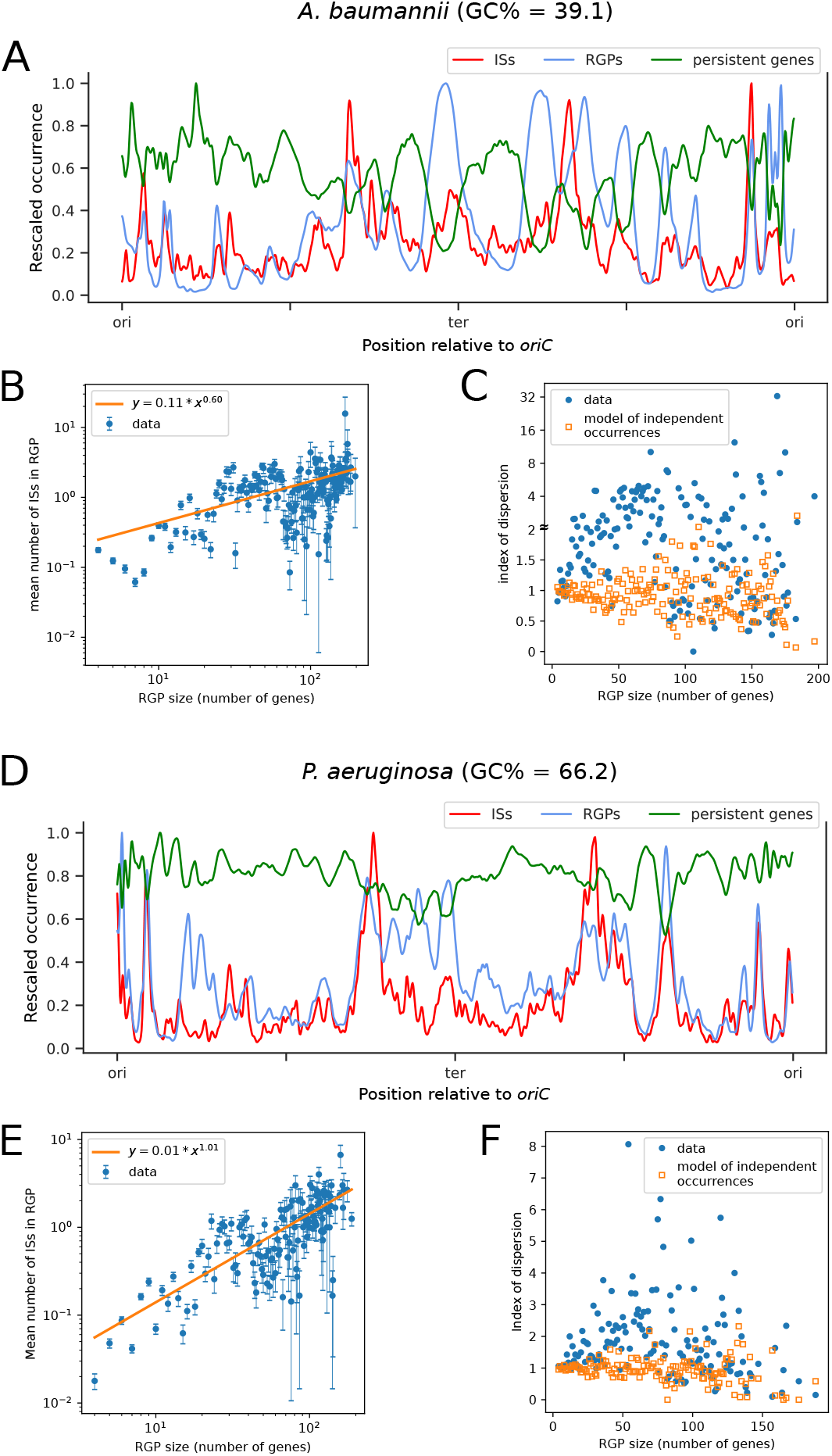
Complementary to Fig. 5. ISs versus RGPs. We show for *A. baumannii* (top) and *P. aeruginosa* (bottom) the same plots as in Fig. 5 obtained in *E. coli*. A,C) Distribution of unique IS positions (red), persistent gene positions (green) and frequency at which a region is a region of genomic plasticity (RGP, blue) across chromosomes. B,D) Mean number of IS copies as a function of RGP size (number of genes within each RGP), computed from all RGPs found in the chromosomes of each species, using a log-log scale – error bars represent the standard error of the mean. The orange line indicates the best power-law fit, revealing as in *E. coli* an almost linear relationship in *P. aeruginosa*. C,E) Index of dispersion (i.e., ratio of the variance to the mean) for the IS content of RGPs as a function of RGP size. The orange symbols represent values obtained in a model in which, within a given RGP, genes have a fixed probability of being an IS, independent of other IS occurrences. These probabilities are determined by the fits in (B,D) and are in average equal to *≃* 1*/*64 for *A. baumannii*, and *≃* 1*/*70 for *P. aeruginosa*.

**Figure S12.**
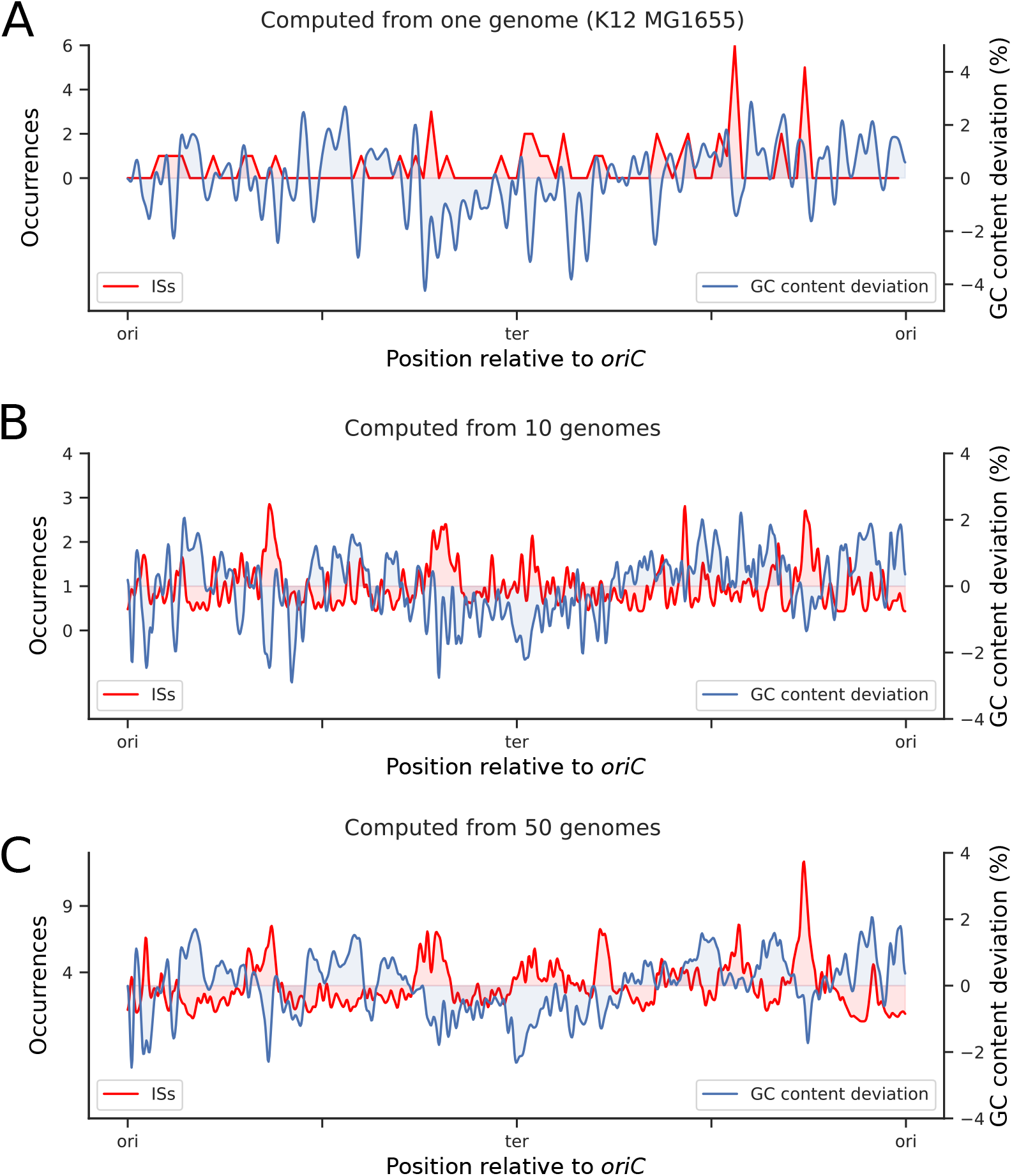
Complementary to Fig. 6. Genomic distribution of ISs versus deviation from average GC content computed by considering different numbers of genomes. A) a single genome (*E. coli* K12 MG1655), B) 10 genomes, and C) 50 genomes randomly selected from a set of 1,917 *E. coli* genomes.

**Figure S13.**
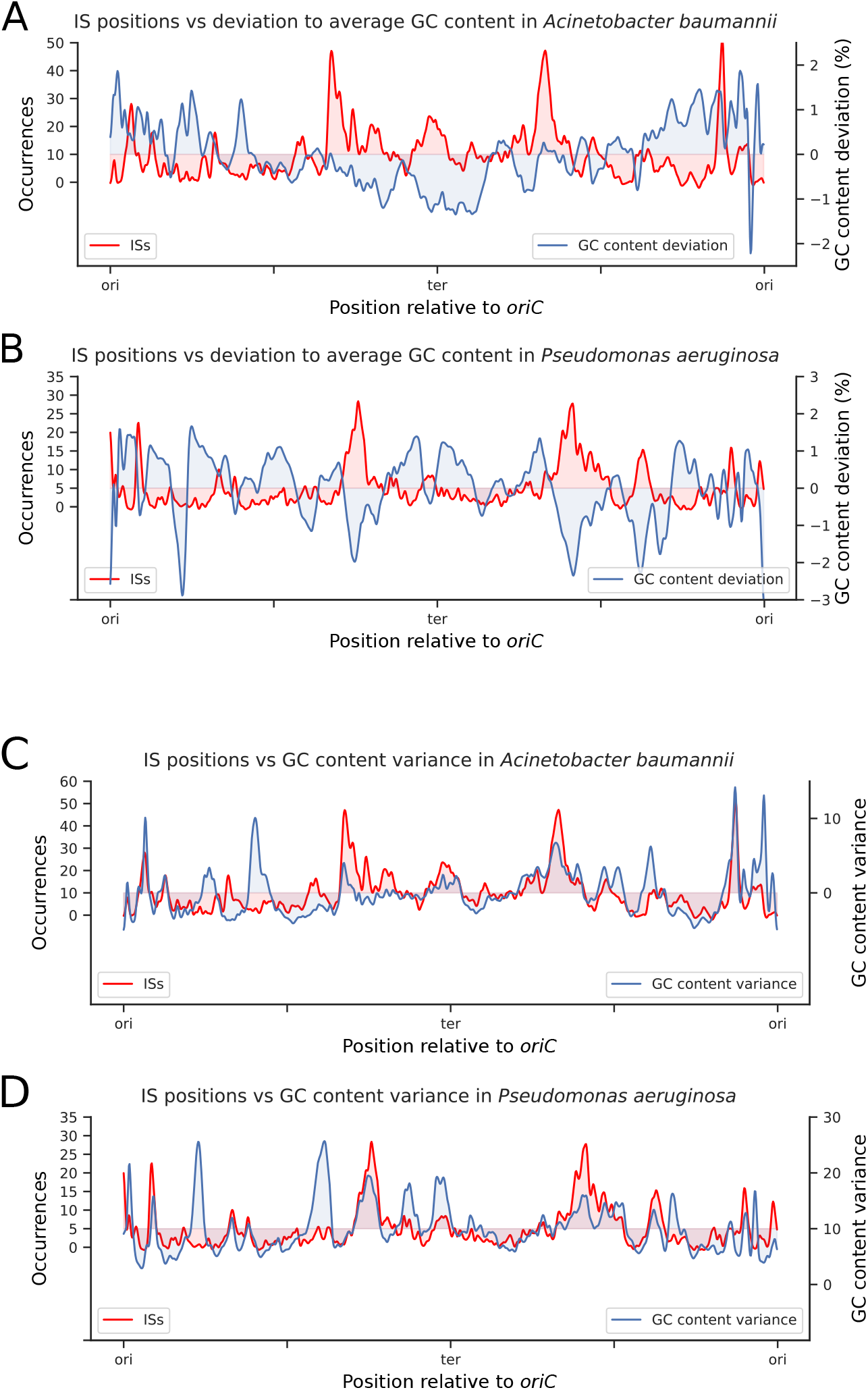
Complementary to Fig. 6. Characterization of the GC content of IS niches in *A. baumannii* and *P. aeruginosa* genomes. A,C) Distribution of unique IS positions (red) and deviation to average GC content (blue) across chromosomes. Pearson correlation: -0.20 for *A. baumannii* and -0.51 for *P. aeruginosa*. B, D) Comparison of IS distribution to the variance in GC content across chromosomes. Pearson correlation: 0.45 for *A. baumannii* and 0.40 for *P. aeruginosa*.

**Figure S14.**
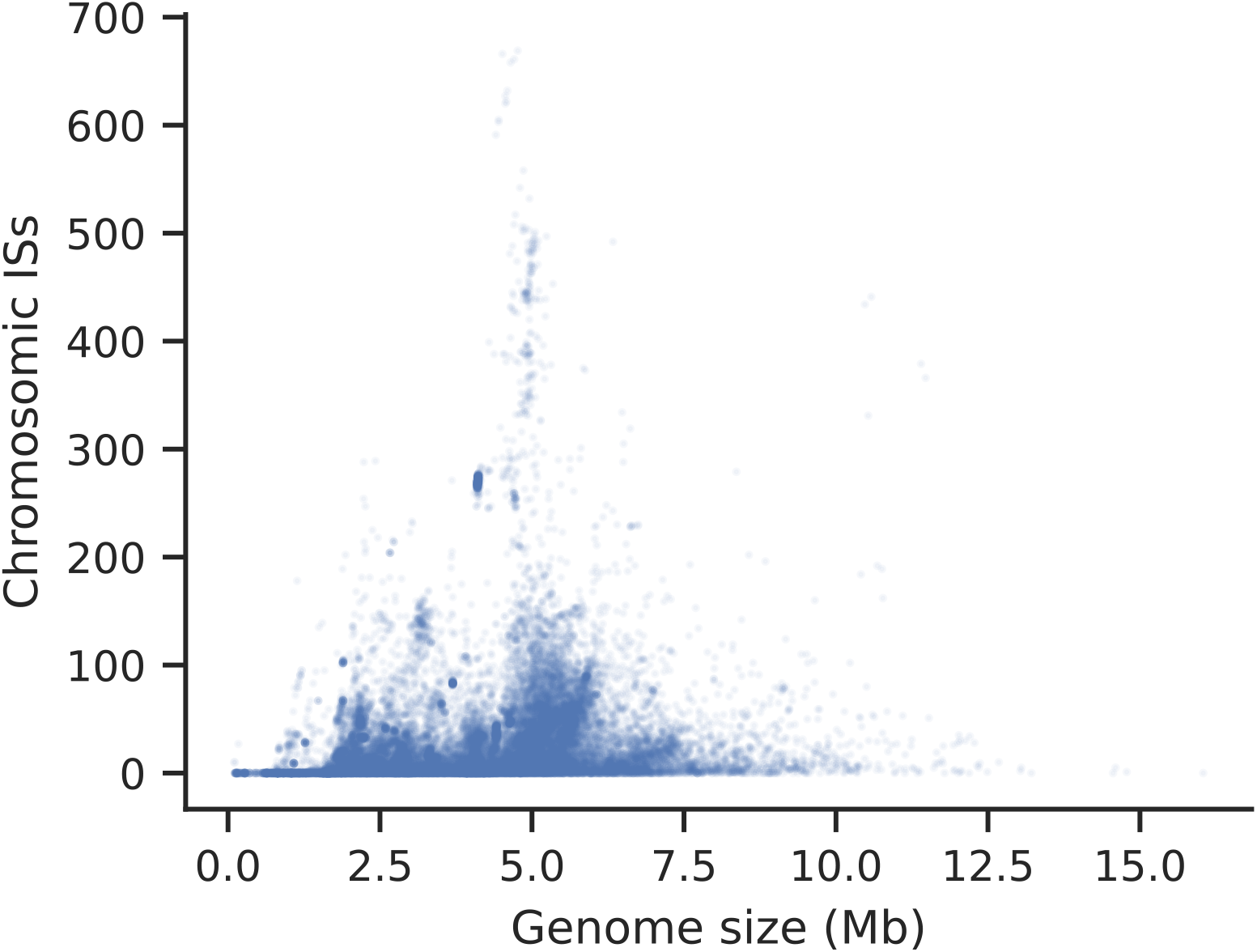
Chromosome size (in Mb) versus total number of ISs found on the chromosome.

**Figure S15.**
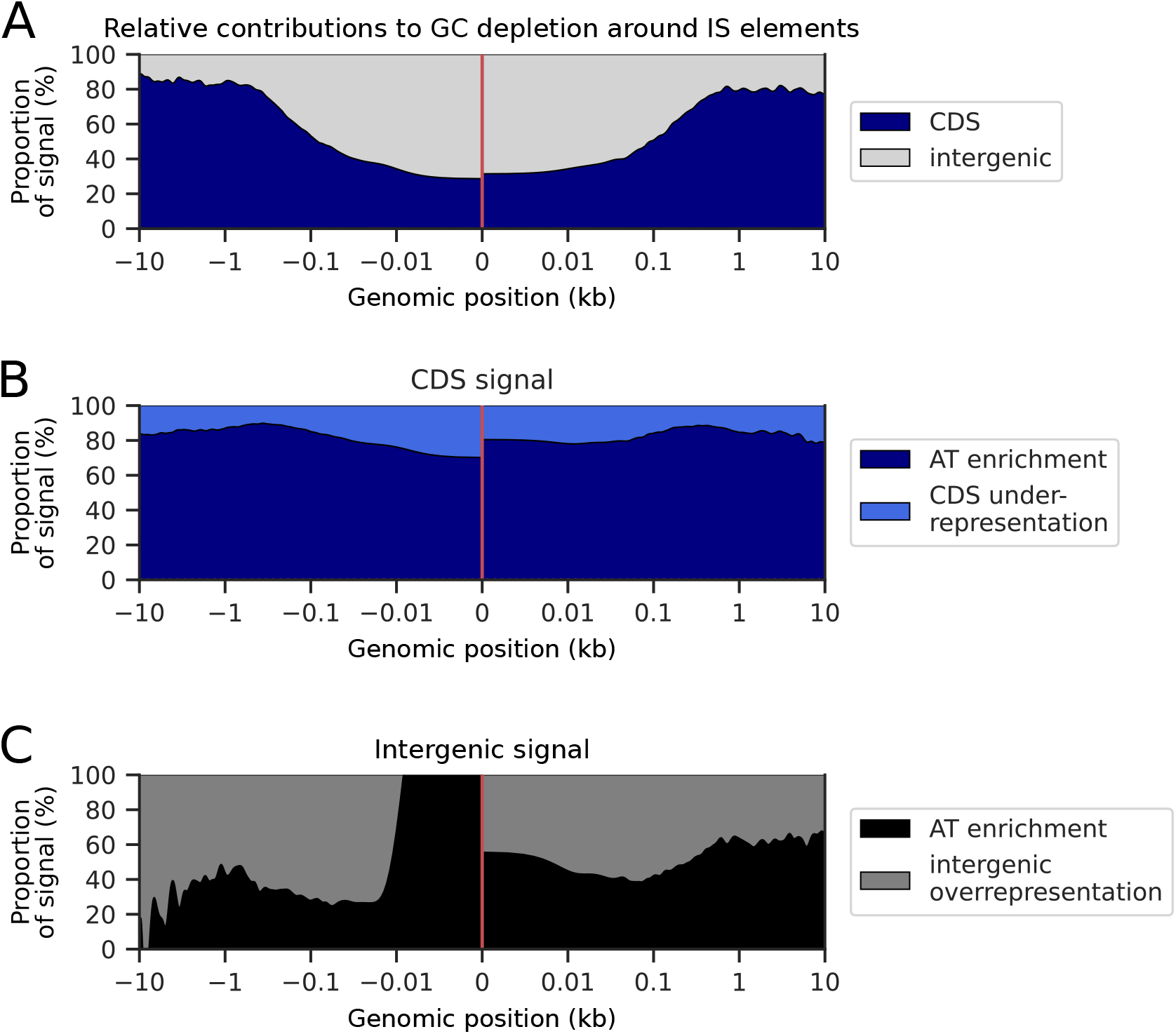
Complementary to Fig. 7. Analysis of the contributions to the GC depletion around ISs. A. Relative contributions of CDSs and intergenic sequences to the GC depletion around ISs. B/C. Contribution of AT enrichment and under/overrepresentation to the AT enrichment signal in CDS/intergenic sequences.

**Figure S16.**
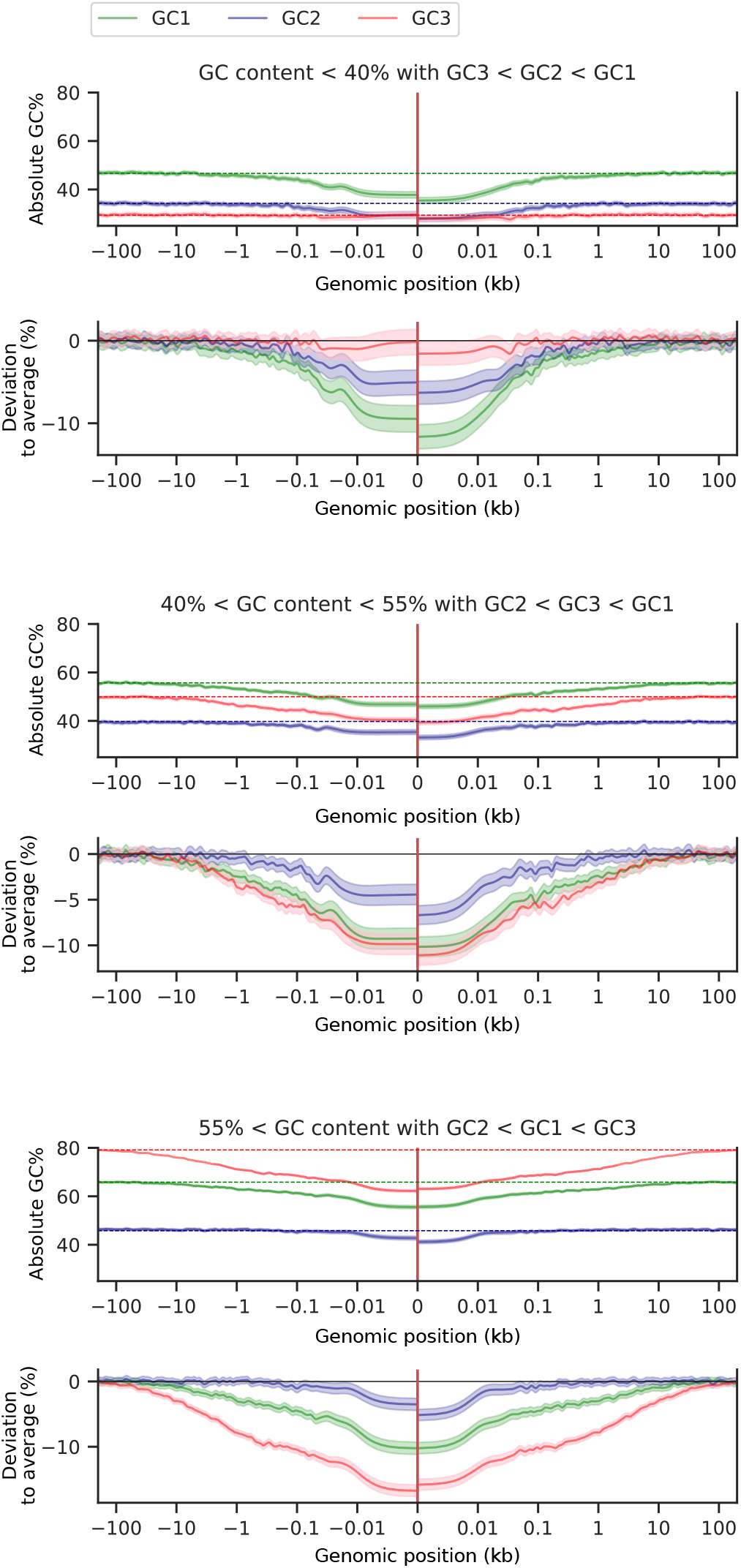
Complementary to Fig. 7. Deviations from average GC content for each codon position. We consider three groups of genomes (68; 69): low, medium, and high GC content, from top to bottom. In each case, the first panel shows the average absolute GC content for each codon position, and the second panel shows the corresponding deviation relative to the average GC content for the associated codon position. Number of genomes: 8,984, 10,542, and 10,973 for low, medium, and high GC content, respectively.

**Figure S17.**
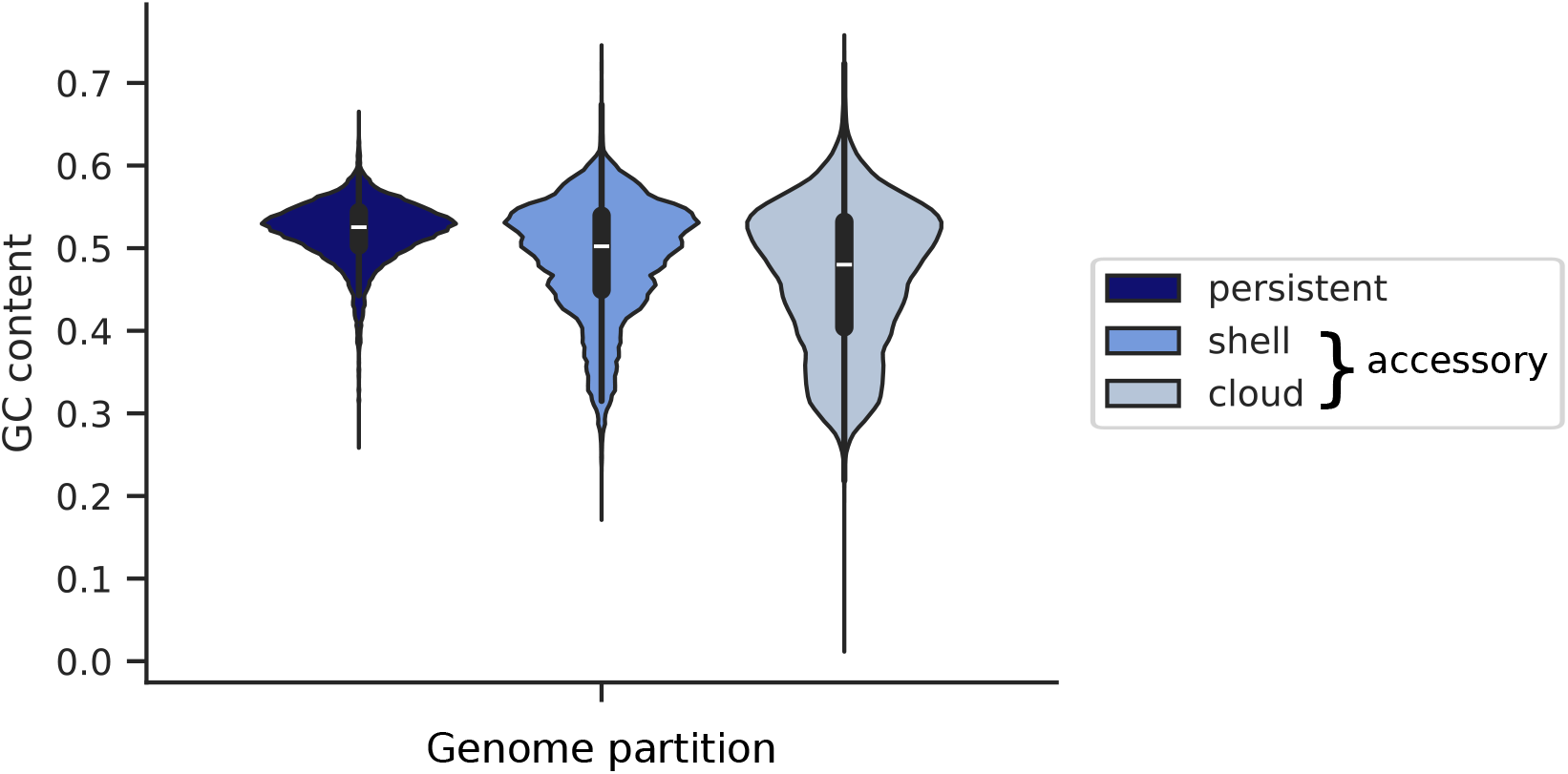
GC content in persistent, shell, and cloud genes in *E. coli* genomes.

**Figure S18.**
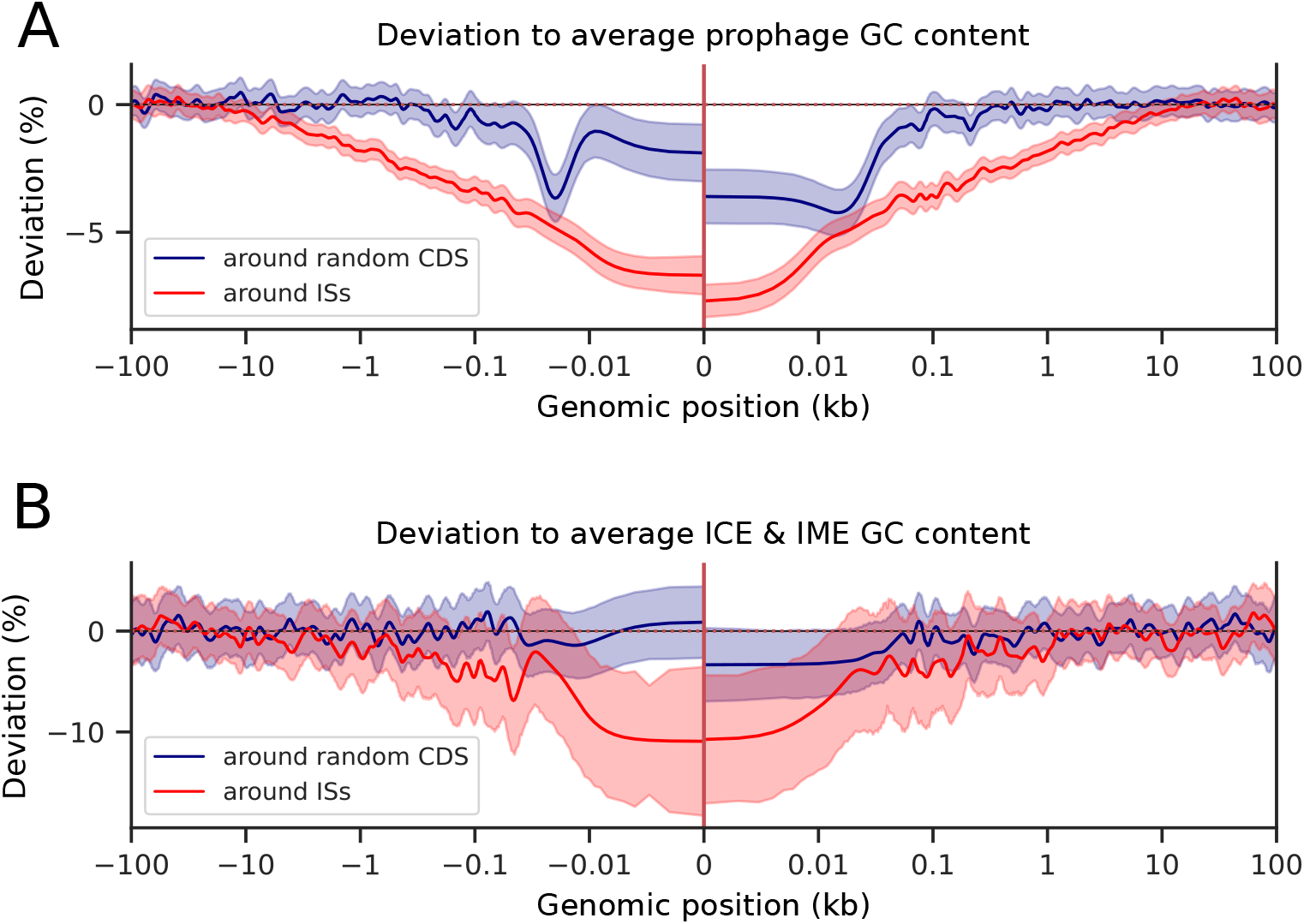
Complementary to Fig. 7. GC content deviation in MGEs around ISs. A. GC content deviation in prophage CDSs around ISs and around random CDSs. B. GC content deviation in ICE and IME sequences around ISs and around random CDSs.

**Figure S19.**
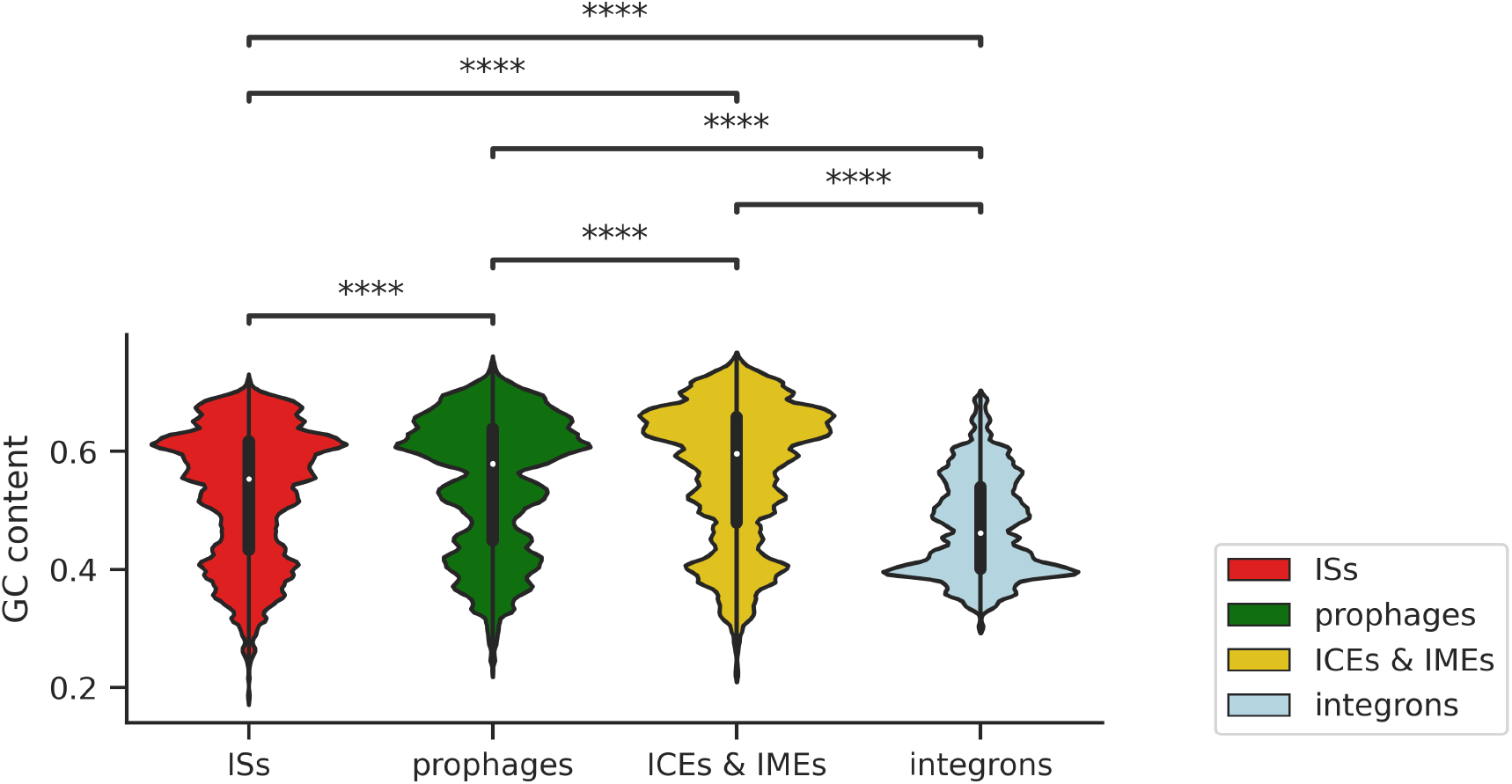
Complementary to Fig. 8. GC content in ISs, prophages, ICEs/IMEs, and integrons. Statistical tests: Mann-Whitney-Wilcoxon test two-sided with Bonferroni correction. All *P* values *P <* 3*e*^*−*308^.

**Figure S20.**
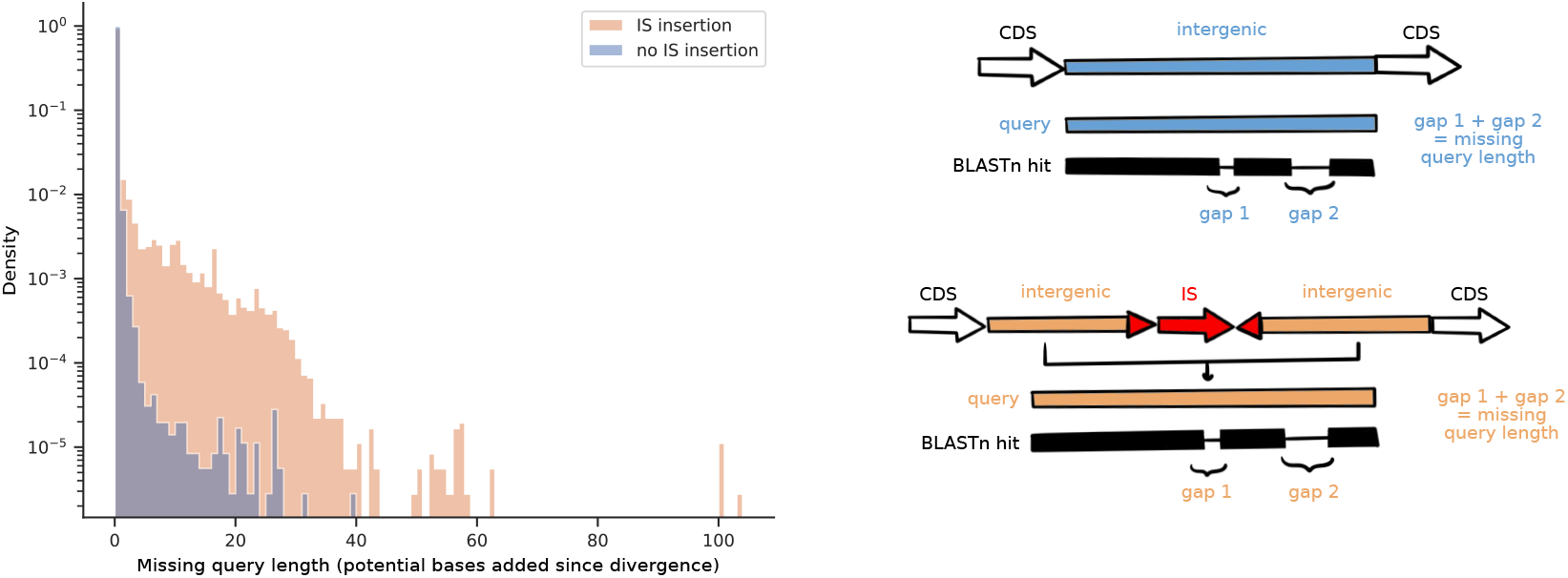
Complementary to Fig. S4A. Analysis of sequences flanking ISs. Red: Missing query length when BLASTing sequences around ISs against the same uninterrupted sequences in other genomes (length of bases present around the IS but not in the sequence found with no IS). Blue: missing query length when BLASTing random sequences with no IS within it against a match in other genomes (control).

## Supplementary Note 5: Supplementary tables

### Supp. Table 1.csv: For each IS family, proportion of copies with a terminator sequence upstream of the ORF inside of the element, upstream of the ORF outside of the element, downstream of the ORF inside of the element, downstream of the ORF outside of the element. For each IS family, a chi2 test was performed to determine if upstream (downstream) terminators were significantly more frequent inside or outside of the element.

### Supp. Table 2.csv: List of COG pairs identified as being in a synteny relationship using a false discovery rate of 0.05. See methodology in (59) for details on the method used to identify these pairs.

